# APIR: Aggregating Universal Proteomics Database Search Algorithms for Peptide Identification with FDR Control

**DOI:** 10.1101/2021.09.08.459494

**Authors:** Yiling Elaine Chen, Xinzhou Ge, Kyla Woyshner, MeiLu McDermott, Antigoni Manousopoulou, Scott B. Ficarro, Jarrod A. Marto, Kexin Li, Leo David Wang, Jingyi Jessica Li

**Affiliations:** Department of Statistics, University of California, Los Angeles, CA 90095, USA; Department of Immuno-Oncology, Beckman Research Institute, City of Hope National Medical Center, Duarte, CA 91010, USA; Department of Quantitative and Computational Biology, University of Southern California, Los Angeles, CA 90089, USA; Department of Cancer Biology and Blais Proteomics Center, Dana-Farber Cancer Institute, Department of Pathology, Brigham and Women’s Hospital and Harvard Medical School, Boston, MA 02215, USA; Department of Pediatrics, City of Hope National Medical Center, Duarte, CA 91010,USA; Interdepartmental Program in Bioinformatics, University of California, Los Angeles, CA 90095, USA; Department of Human Genetics, University of California, Los Angeles, CA 90095, USA; Department of Computational Medicine, University of California, Los Angeles, CA 90095, USA; Department of Biostatistics, University of California, Los Angeles, CA 90095, USA, USA

**Author notes:** Equal contribution (Yiling Elaine Chen, Xinzhou Ge, Kyla Woyshner, MeiLu McDermott). Corresponding authors: (Li JJ), (Wang LD).

**Keywords:** Shotgun proteomics, Peptide-spectrum match, Peptide identification, Aggregation of lists, FDR control

## Abstract

Advances in mass spectrometry (MS) have enabled high-throughput analysis of proteomes in biological systems. The state-of-the-art MS data analysis relies on database search algorithms to quantify proteins by identifying peptide-spectrum matches (PSMs), which convert mass spectra to peptide sequences. Different database search algorithms use distinct search strategies and thus may identify unique PSMs. However, no existing approaches can aggregate all user-specified database search algorithms with a guaranteed increase in the number of identified peptides and control on the false discovery rate (FDR). To fill in this gap, we propose a statistical framework, Aggregation of Peptide Identification Results (APIR), that is universally compatible with all database search algorithms. Notably, under an FDR threshold, APIR is guaranteed to identify at least as many, if not more, peptides as individual database search algorithms do. Evaluation of APIR on a complex proteomics standard shows that APIR outpowers individual database search algorithms and empirically controls the FDR. Real data studies show that APIR can identify disease-related proteins and post-translational modifications missed by some individual database search algorithms. The APIR framework is easily extendable to aggregating discoveries made by multiple algorithms in other high-throughput biomedical data analysis, e.g., differential gene expression analysis on RNA sequencing data. The APIR R package is available at https://github.com/yiling0210/APIR.

## Introduction

Proteomics studies have discovered essential roles of proteins in complex diseases such as neurodegenerative disease [1] and cancer [2, 3]. These studies have demonstrated the potential of using proteomics to identify clinical biomarkers for disease diagnosis and therapeutic targets for disease treatment. In recent years, proteomics analytical technologies, particularly tandem mass spectrometry (MS)-based shotgun proteomics, have advanced immensely, thus enabling high-throughput identification and quantification of proteins in biological samples. Compared to prior technologies, shotgun proteomics has simplified sample preparation and protein separation, reduced time and cost, and saved procedures that may result in sample degradation and loss [4]. In a typical shotgun proteomics experiment, a protein mixture is first enzymatically digested into peptides, i.e., short amino acid chains up to approximately 40-residue long; the resulting peptide mixture is then separated and measured by tandem MS into tens of thousands of mass spectra. Each mass spectrum encodes the chemical composition of a peptide; thus, the spectrum can be used to identify the peptide’s amino acid sequence and post-translational modifications, as well as to quantify the peptide’s abundance with additional weight information (**Figure 1**A).

**Figure 1.**
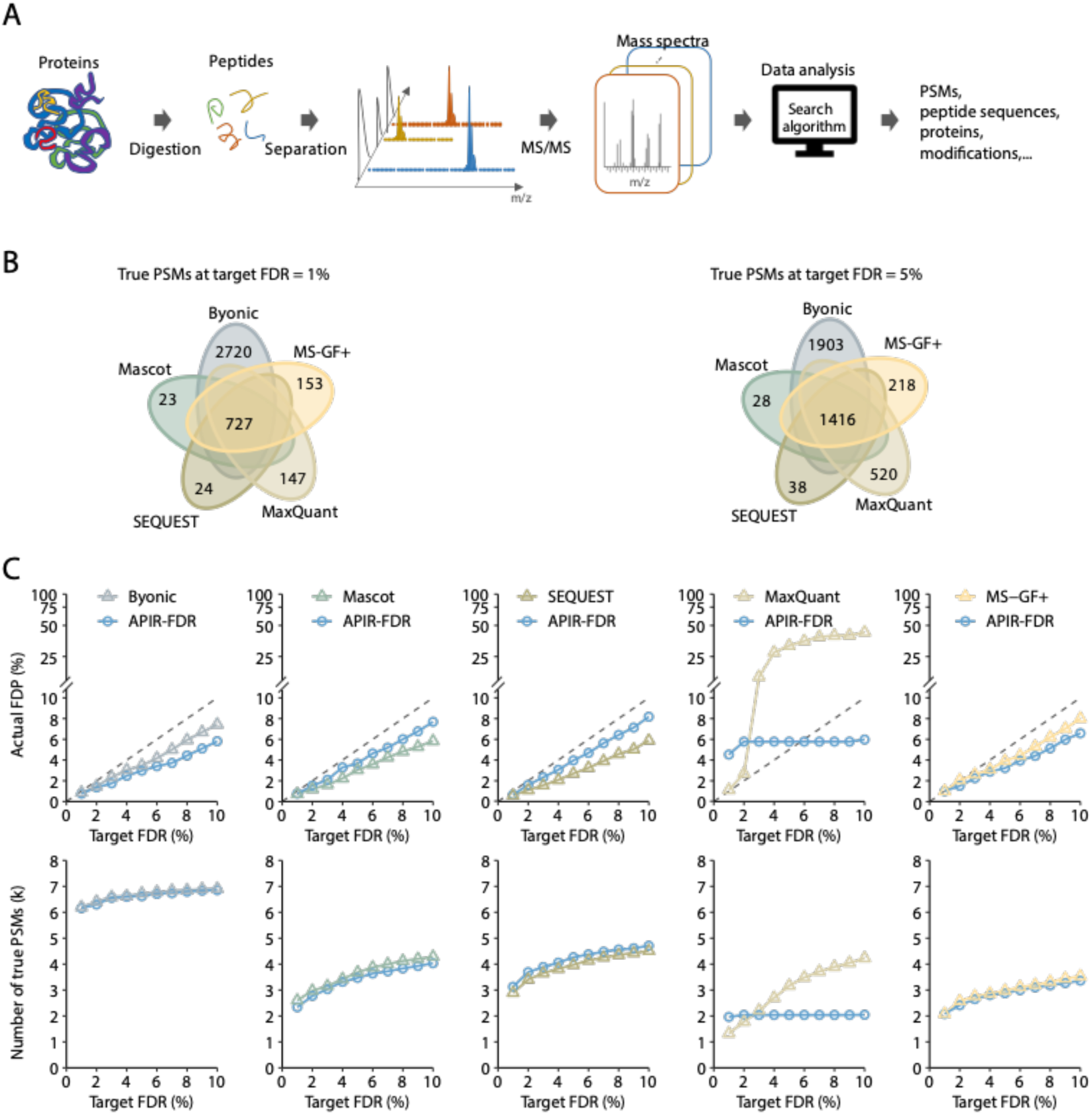
The workflow of shotgun proteomics and benchmarking search algorithms on proteomics standard. **A**. The workflow of a typical shotgun proteomics experiment. The protein mixture is first enzymatically digested into peptides, i.e., short amino acid chains up to approximately 40-residue long; the resulting peptide mixture is then separated and measured by tandem MS into tens of thousands of mass spectra. Each mass spectrum encodes the chemical composition of a peptide. Then a database search algorithm is used to identify the peptide’s amino acid sequence and post-translational modifications, as well as to quantify the peptide’s abundance. **B**. Venn diagrams showing the overlap of true PSMs identified by the five database search algorithms from the *Pfu* proteomics standard dataset under the FDR threshold *q* = 1% (left) or *q* = 5% (right). **C**. The FDP and power of each database search algorithm on the *Pfu* proteomics standard dataset at the FDR threshold *q* ∈ {1%, …, 10%}. MS, mass spectrometry; PSM, peptide-spectrum matches; FDR, false discovery rate; FDP, false discovery proportion; APIR, Aggregation of Peptide Identification Results.

Since the development of shotgun proteomics, numerous database search algorithms have been developed to automatically convert mass spectra into peptide sequences. Popular database search algorithms include SEQUEST[5], Mascot [6], MaxQuant[7], Byonic [8], and MS-GF+ [9], among many others. A database search algorithm takes as input the mass spectra from a shotgun proteomics experiment and a protein database (called the “target database”) that contains known protein sequences (called “target sequences”). For each mass spectrum, the algorithm identifies the best matching peptide sequence, i.e., a subsequence of a protein sequence, from the database; we call this process “peptide identification,” whose result is a “peptide-spectrum match” (PSM). However, due to data imperfection (such as low-quality mass spectra, data processing mistakes, and protein database incompleteness), the identified PSMs often consist of many false PSMs, causing issues in the downstream system-wide identification and quantification of proteins [10].

To ensure the accuracy of PSMs, the false discovery rate (FDR) has been used as the most popular statistical criterion [11–18]. Technically, the FDR is defined as the expected proportion of false PSMs among the identified PSMs; in other words, a small FDR indicates good accuracy of PSMs. To control the FDR, the standard approach is the target-decoy search, which utilizes a “decoy database” consisting of known, non-existent protein sequences (called “decoy sequences”) [10]. Two common strategies for target-decoy search are concatenated search and parallel search. The concatenated search strategy finds the best match of a mass spectrum in a concatenated database containing both target sequences and decoy sequences; hence, the match (i.e., PSM) corresponds to either a target sequence or a decoy sequence. In contrast, the parallel search strategy finds the best match of a mass spectrum in the target database and the decoy database separately; hence, the spectrum has two best matches, one with a target sequence (i.e., a target PSM) and the other with a decoy sequence (i.e., a decoy PSM). Based on the target-decoy search results (regardless of being concatenated or parallel), including target PSMs and decoy PSMs with matching scores, multiple procedures that are p-value-based or p-value-free have been proposed to control the FDR of a database search algorithm’s identified target PSMs [14][19–21].

However, controlling the FDR is only one side of the story. Because shotgun proteomics experiments are costly, a common goal of database search algorithms is to identify as many true PSMs as possible to maximize the experimental output, in other words, to maximize the identification power given a target, user-specified FDR threshold (e.g., 1% or 5%).

It has been observed that, with the same input mass spectra and FDR threshold, different database search algorithms often find largely distinct sets of PSMs [22–26]. In this study, we confirmed this observation using our in-house dataset, the first publicly available complex proteomics standard dataset from *Pyrococcus Furiosus* (*Pfu*) that approximates the dynamic range of a typical proteomics experiment. We first benchmarked five popular database search algorithms---Byonic [8], Mascot [6], SEQUEST[5], MaxQuant [7], and MS-GF+ [9]---on the proteomics standard dataset using an FDR assessment approach similar to that in [27]. Our results confirmed that these five algorithms were designed to capture unique sets of PSMs (see Results for details). Hence, it is reasonable to use aggregation methods to combine individual database search algorithms’ outputs to boost the power of identifying peptides from shotgun proteomics data.

In the proteomics field, existing aggregation methods include Scaffold [25], MSblender [18], FDRAnalysis [28], iProphet [17], ConsensusID [16], PepArML [11], and a multi-stage method by Ning et al. [29]. Among these seven methods, except FDRAnalysis, which has been shown infeasible for high-throughput proteomics [22], the rest have at least one of the two major drawbacks: (1) limited compatibility with database search algorithms and (2) lack of guarantee for identifying more peptides under the same FDR threshold. For the first drawback, except ConsensusID, the other six aggregation methods unanimously limit the choices of database search algorithms. As for the second drawback, although empirical evidence shows that, on some datasets, these aggregation methods, except the multi-stage method by Ning et al. [29], may identify more peptides than those identified by individual database search algorithms, none of these aggregation methods is guaranteed to do so by algorithm design.

In addition to the above aggregation methods developed for proteomics data, generic statistical methods developed for aggregating rank lists are in theory applicable to aggregating the PSM lists output by database search algorithms. However, none of these generic methods have been developed into software packages compatible with database search algorithms, nor are they guaranteed to identify more peptides given an FDR threshold (many generic methods aggregate rank lists without FDR control). Therefore, the field calls for a robust, powerful, and flexible aggregation method that allows researchers to reap the benefits of the diverse and ever-growing database search algorithms.

Here we propose Aggregate Peptide Identification Results (APIR), a statistical framework that aggregates peptide identification results from multiple database search algorithms with FDR control. Compared to the existing aggregation methods, APIR offers the following three advantages simultaneously: first, APIR is open-source and universally adaptive to database search algorithms that output PSMs with matching scores (e.g., q-values or posterior error probabilities); second, APIR is guaranteed to identify at least as many as, if not more, peptides than individual database search algorithms do; third, APIR empirically controls the FDR in simulation and real-data benchmark studies. Hence, APIR is a robust, flexible framework that enhances the power while controlling the FDR of peptide identification from shotgun proteomics data.

Note that the framework of APIR could be easily extended to aggregate discoveries made by multiple algorithms in other high-throughput biomedical data analysis, such as differential gene expression analysis on RNA sequencing data.

## Method

We propose APIR to aggregate multiple database search algorithms’ output PSMs. Designed to control the FDR of aggregated PSMs, APIR is a sequential framework applied to individual database search algorithms’ output PSMs. To benchmark APIR and existing database search algorithms, we also generated the first publicly available complex proteomics standard from *Pfu* to approximate the dynamic range of a typical proteomics experiment. Below we first introduce the methodology of APIR, including APIR-FDR and the sequential framework for aggregating PSMs. Then we introduce the experimental details on how we generated the proteomics standard dataset and used it for benchmarking purposes.

### APIR methodology

Aside from a user-specified FDR threshold *q* (e.g., 5%), APIR takes as input the target-decoy search results from the database search algorithms users would like aggregate [10]. Specifically, APIR requires from each database search algorithm a list of target PSMs with matching scores and a list of decoy PSMs with matching scores. To maximize power, we recommend users to extract the entire lists of target PSMs and decoy PSMs by setting the internal FDR of each database search algorithm to 100%. Note that the target-decoy search strategy referred to herein does not include the FDR estimation procedure criticized by [19].

To facilitate downstream analysis, APIR also reports the master protein, the post-translational modifications, and the abundance of each identified PSM, if applicable. See File S1 for details on these post-processing steps.

#### APIR-FDR: FDR control on any individual search algorithm

The core component of APIR is APIR-FDR, an umbrella FDR-control procedure for each individual database search algorithm’s identified target PSMs. APIR-FDR takes as input an FDR threshold *q*, a list of target PSMs with matching scores, and a list of decoy PSMs with matching scores. APIR-FDR then outputs the identified target PSMs. As an umbrella FDR-control procedure, APIR-FDR can be p-value-based or p-value-free, including all possible procedures that can control the FDR for an individual database search algorithm’s identified target PSMs. Below we describe three exemplar options for APIR-FDR: a p-value-based option and two p-value-free options.

To facilitate our discussion, we introduce some notations. Let *m* and *n* denote the numbers of target PSMs and decoy PSMs, respectively, outputted by a database search algorithm. We denote the matching scores of target PSMs and decoy PSMs as *T*_1_, …, *T_m_* and *D*_1_, …, *D_n_*, respectively. Without loss of generality, we assume that the matching scores are positive, and a larger matching score indicates a higher chance for a PSM to be a true match. For instance, if the output of a database search algorithm contains PSMs with q-values or e-values (whose smaller values indicate more likely true matches), then we would define the negative log-transformed q-values or e-values as matching scores.

First, a p-value-based FDR-control procedure applies to both concatenated and parallel target-decoy search strategies [14]. It assumes that the matching scores of decoy PSMs and false target PSMs are independently and identically distributed, and it constructs a null distribution by pooling the matching scores of decoy PSMs *D*_1_, …, *D_n_*. Then, it computes a p-value for the *i*-th target PSM as the tail probability right of *T_i_*, i.e., 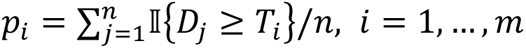. Given the FDR threshold *q* and the p-values *p*_1_, …, *p_m_*, this procedure applies the Benjamini-Hochberg procedure [30] to set a p-value threshold *p*_thre_ (*q*) and outputs {*i* = 1, …, *m*: *p_i_* ≤ *p*_thre_ (*q*)} as the indices of the identified target PSMs.

Second, a p-value-free FDR-control procedure (as a clarification, this procedure was referred to as the target-decoy search strategy in [31], different from the terminologies we use) also applies to both concatenated and parallel search strategies. When used with concatenated search results, for a given matching score *x*, this procedure counts the numbers of target PSMs and decoy PSMs with matching scores at least *x* as 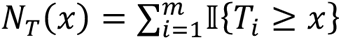 and 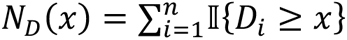 respectively. This procedure then estimates the FDR of target PSMs with matching scores at least *x* as 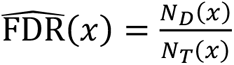 When used with parallel search results, the procedure needs to estimate π_1_, the proportion of false PSMs among the target PSMs. This proportion is unknown but can be conservatively estimated. This is done by examining PSMs with scores near zero and then calculating the ratio of the number of decoy PSMs to the number of target PSMs in this subset of PSMs, as outlined in reference [14]. With the estimated 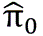, the procedure then estimates the FDR of target PSMs with matching scores at least *x* as 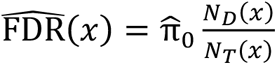. Given the FDR threshold *q*, this procedure outputs 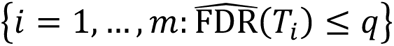 as the indices of the identified target PSMs.

Third, an alternative p-value-free FDR-control procedure is Clipper, which works for parallel search results by design (Clipper controls the FDR by contrasting two conditions, which correspond to a mass spectrum’s target match and decoy match in parallel search) [10]. In parallel search results, we assume that the first *s* ≤ min(*m*, *n*) target PSMs can be paired one-to-one with decoy PSMs. Then we arrange the decoy PSM indices in the way that the *i*-th decoy PSM shares the same mass spectrum with the *i*-th target PSM for 1 ≤ *i* ≤ *s* (note that *s*/*m* is close to 1 in most parallel search results). Clipper first constructs a contrast score *C_i_* = *T_i_* − *D_i_* for *i* = 1, …, *s*; note that the contrast score may be defined in other forms [32]. Then given the FDR threshold *q*, Clipper finds a contrast score cutoff 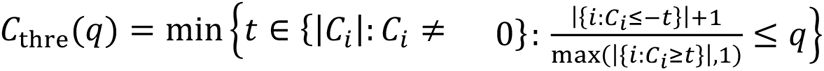, and outputs {*i* = 1, …, *s*: *C_i_* ≥ *C*_thre_ (*q*)} as the indices of the identified target PSMs, where |{*i*: *C*_*i*_ ≤ −*t*}| indicates the number of *C*_1_, … *C*_s_ that are no greater than −*t*. For the *i*-th target PSM, *i* = 1, …, *s*, Clipper estimates its FDR as 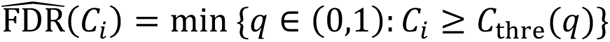. In comparison to the p-value-free procedure outlined previously, Clipper is more conservative (owing to the “+1” in the numerator) and flexible. Notably, Clipper is similar to the above p-value-free FDR-control procedure if the contrast score is defined as *C_i_* = max(*T_*i*_*, *D_*i*_*). Compared to the p-value-free FDR-control procedure outlined in the previous paragraph, Clipper has three advantages: (1) Clipper does not require the estimation of π_0_ in parallel search; (2) Clipper’s estimated FDR 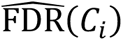 monotonically decreases as the contrast score *C_*i*_* increases, resulting in better power in numerous instances; (3) Clipper is more flexible because its contrast score *C_*i*_* can be defined in various ways.

The FDR-control procedures described above are just three examples. Any other procedures that control the FDR can also be used as options for APIR-FDR.

#### APIR: a sequential framework for aggregating multiple search algorithms’ identified target PSMs with FDR control

Given the FDR threshold *q* and multiple database search algorithms’ outputs (including all target PSMs and decoy PSMs with matching scores), the sequential framework of APIR identifies target PSMs (with the FDR controlled under *q*) by combining these database search algorithms’ outputs based on a mathematical fact: if disjoint sets of discoveries all have the false discovery proportion (FDP; also known as the empirical FDR) under *q*, then their union set also has the FDP under *q*. Hence, the sequential framework of APIR is designed to find disjoint sets of target PSMs from database search algorithms’ outputs. The final output of APIR is the union of these disjoint sets, which is guaranteed to contain more unique peptides than what could be identified by a single database search algorithm.

Suppose we are interested in aggregating *K* algorithms. Accordingly, the sequential approach will consist of a maximum of *K* rounds. Let *W*_A_ denote the set of target PSMs output by the *k*-th algorithm, *k* = 1, …, *K*. In Round 1, APIR applies APIR-FDR to each algorithm’s output with the FDR threshold *q*. Denote the identified target PSMs from the *k*-th algorithm by *U*_1*k*_ ⊂ *W_k_*. Define *J*_1_ ∈ {1, …, *K*} to be the algorithm index such that *U*_1*J*1_ contains the largest number of unique peptides among *U*_11_, …, *U*_1*k*_. We use the number of unique peptides rather than the number of PSMs because peptides are more biologically relevant than PSMs. In Round 2, APIR first excludes all target PSMs output by algorithm *J*_1_, identified or unidentified in Round 1, i.e., *W_J_*_1_, from the outputs of the remaining database search algorithms, resulting in reduced sets of candidate target PSMs *W*_1_ ∖ *W_J_*_1_ …, *W_k_* ∖ *W_J_*_1_. Then APIR applies APIR-FDR with FDR threshold *q* to these reduced sets except *W_J_*_1_ ∖ *W_J_*_1_ = ∅. Denote the resulting sets of identified target PSMs by *U*_2*k*_ ⊂ (*W_k_* ∖ *W_J_*_1_), *k* ∈ {1, …, *K*} ∖ {*J*_1_}. Again APIR finds algorithm *J*_2_ such that *U*_2*J*2_ contains the largest number of unique peptides. APIR repeats this procedure in the subsequent rounds. Specifically, in Round ℓ with ℓ ≥ 2, APIR first excludes all target PSMs output by the selected ℓ − 1 algorithms from the outputs of remaining database search algorithms and applies APIR-FDR. That is, APIR applies APIR-FDR with FDR threshold *q* to identify a set of identified PSMs *U*_ℓk_ from *W_k_* ∖ (*W_J_*_1_ ∪ ⋯ ∪ *W_J_*_ℓ-1_), the reduced candidate pool of algorithm *k* after the previous ℓ rounds, for algorithms *k* ∈ {1, …, *K*} ∖ {*J*_1_, …, *J*_ℓ-1_}. Then APIR finds the algorithm, which we denote by *J*_ℓ_, such that *U_ℓjℓ_* contains the largest number of unique peptides. Finally, APIR outputs *U*_1*j*1_ ∪ ⋯ ∪ *U*_k*j*k_ as the set of identified target PSMs. By adopting this sequential approach, APIR is guaranteed to identify at least as many, if not more, unique peptides as those identified by a single database search algorithm; under the assumption that APIR-FDR controls the FDR for each algorithm’s identified target PSMs, APIR controls the FDR of the identified target PSMs under *q*, APIR controls the FDR of the identified target PSMs under *q*. See **Figure 2** for an illustration of APIR’s sequential approach.

**Figure 2.**
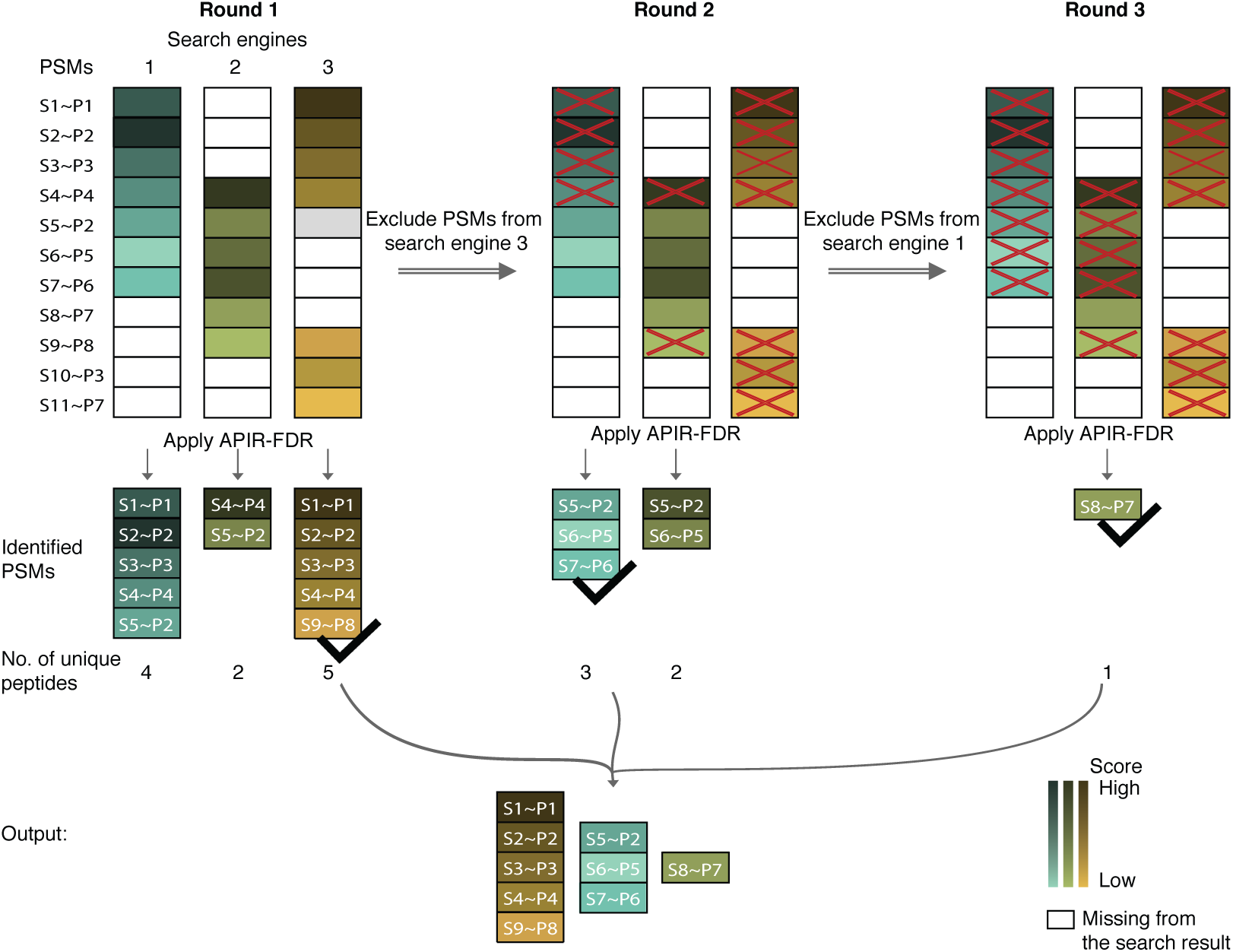
Illustration of APIR for aggregating three database search algorithms. We use S1∼P1 to denote a PSM of mass spectrum S1 matched to peptide sequence P1. In the output of a database search algorithm, a PSM with a higher matching score is marked by a darker color. White boxes indicate PSMs missing from the output. APIR adopts a sequential approach to aggregate the three database search algorithms. In Round 1, APIR applies APIR-FDR to identify a set of target PSMs from the output of each database search algorithm. APIR then selects the algorithm whose identified PSMs contain the largest number of unique peptides, and the identified PSMs are considered identified by APIR. In this example, APIR identified the same number of PSMs from algorithms 1 and 3 but more unique peptides from algorithm 3; hence, APIR selects algorithm 3. In Round 2, APIR excludes all PSMs, either identified or unidentified by the selected database search algorithm in Round 1 (algorithm 3 in this example), from the output of the remaining database search algorithms. Then APIR applies APIR-FDR to find the algorithm whose identified PSMs contain the largest number of unique peptides (algorithm 1 in this example). APIR repeats Round 2 in the subsequent rounds until all database search algorithms are selected. Finally, APIR outputs the union of the PSMs identified in each round. PEP, posterior error probability.

Notably, APIR specifically controls the FDR of identified target PSMs so it excludes the identified target PSMs, instead of spectra, of one algorithm in each step. In instances where different algorithms match the same spectrum to distinct peptides, APIR may identify both PSMs as valid discoveries. While at least one of these two PSMs is a false discovery, the overall FDR for the identified PSMs remains controlled under this framework.

#### Complex proteomics standard dataset generation

We describe the experimental details of running the tandem MS analysis on a *Pfu* proteomics standard sample. The complex proteomics standard (CPS) (part number 400510) was purchased from Agilent (Agilent, Santa Clara, CA, USA). CPS contains soluble proteins extracted from the archaeon *Pfu*. All other chemicals were purchased from Sigma Aldrich (Sigma Aldrich, St. Louis, MO, USA). The fully sequenced genome of *Pfu* encodes for approximately 2000 proteins that cover a wide range of size, pI, concentration level, hydrophobic/hydrophilic character, etc. CPS (500 μg total protein) was dissolved in 100 μL of 0.5 M triethylammonium bicarbonate (TEAB) and 0.05% sodium dodecyl sulfate (SDS) solution. Proteins were reduced using tris(2-carboxyethyl)phosphine hydrochloride (TCEP) (4 μL of 50 mM solution added to the protein mixture and sample incubated at 60℃ for 1 hour) and alkylated using methyl methanethiosulfonate (MMTS) (2 μL of 50 mM solution added to the protein mixture and sample incubated at room temperature for 15 minutes). 20 μg trypsin dissolved 1:1 in ultrapure water was added to the sample, and this was incubated overnight (16 hours) in the dark at 37℃ to enzymatically digest the proteins. The tryptic peptides were cleaned with C-18 tips (part number 87784) from Thermo Fisher Scientific (Thermo Fisher Scientific, Waltham, MA, USA) following the manufacturer’s instructions. Peptides were LC-MS analyzed using the Ultimate 3000 uPLC system (EASY-Spray column, part number ES803A, Thermo Fisher Scientific) hyphenated with the Orbitrap Fusion Lumos mass spectrometry instrument (Thermo Fisher Scientific). Peptides were fragmented using low-energy CID and detected with the linear ion trap detector.

After running tandem MS analysis, we obtained 49,303 mass spectra from *Pfu*. We then adopted an approach similar to that in [27] for benchmarking database search algorithms and aggregation methods. Specifically, we first constructed a target database by concatenating the *Pfu* database, the Uniprot Human database [33], and two contaminant databases: the CRAPome [34] and the contaminant databases from MaxQuant. In the target database construction, we removed human proteins that contain *Pfu* peptides (via *in silico* trypsin digestion). Contaminant databases consist of sequences commonly identified as contaminants in MS experiments. Given that PSMs resulting from unintended sources, such as contamination, are unavoidable in MS experiments, PSMs originating from both the Pfu database and contaminant databases are considered as true PSMs. Conversely, PSMs from the human database, after excluding all Pfu proteins, are cconsidered as false PSMs. Finally, we input the 49,303 mass spectra and the reference database into database search algorithms. To evaluate a database search algorithm or an aggregation method, we consider its output PSMs, peptides, and proteins as true if and only if they belong to either *Pfu* or the two contaminant databases. The in silico digestion was done to take out any human proteins that contain peptides that could also be derived from Pfu. The *in silico* digestion was performed in Python using the pyteomics.parser function from pyteomics with the following settings: Trypsin digestion, two allowed missed cleavages, minimum peptide length of six amino acid residues [35, 36].

## Results

To verify the motivation and demonstrate the advantages of APIR, we conducted simulation and real data studies. First, we benchmarked five popular database search algorithms---Byonic, Mascot, SEQUEST, MaxQuant, and MS-GF+ — coupled with APIR-FDR options (p-value-based or p-value-free) on our *Pfu* proteomics standard dataset. Second, we designed simulation studies to benchmark APIR against two naïve aggregation approaches: intersection and union of different database search algorithms’ identified PSM sets. Third, to demonstrate the power of APIR, we applied it to five real datasets, including our proteomics standard dataset, three acute myeloid leukemia (AML) datasets, and a triple-negative breast cancer (TNBC) dataset. Notably, we generated two of the three AML datasets from bone marrow samples of AML patients with either enriched or depleted leukemia stem cells (LSCs) for studying the disease mechanisms of AML. Finally, we investigated and verified additional proteins found by APIR and performed differentially expressed peptide analysis on the APIR results.

Although we focused on five database search algorithms, APIR is universally applicable to other database search algorithms such as MSFragger [37] and Open-pFind [38]. Because nearly all database search algorithms output q-values or posterior error probabilities (PEPs) of PSMs, we used – log_10_ –transformed PEPs from MaxQuant and − log_!1_ –transformed q-values from the other four database search algorithms as PSMs’ matching scores to demonstrate the wide applicability of APIR.

### Benchmarking five database search algorithms on the *Pfu* proteomics standard

We first benchmarked five popular database search algorithms---Byonic, Mascot, SEQUEST, MaxQuant, and MS-GF+---on the *Pfu* proteomics standard dataset. Our evaluation results in Figure 1B show that the five individual database search algorithms indeed capture unique true PSMs in this proteomics standard dataset at FDR thresholds *q* = 1% and 5%. Notably, at *q* = 1%, the number of true PSMs identified by Byonic alone (2720) is nearly four times the number of true PSMs identified by all five algorithms (727). At *q* = 5%, Byonic again identifies more unique true PSMs (1903) than the true PSMs identified by all five algorithms (1416). Moreover, MaxQuant and MS-GF+ also demonstrate distinctive advantages: MaxQuant identifies 147 and 520 unique true PSMs, while MS-GF+ identifies 153 and 218 unique true PSMs at *q* = 1% and 5%, respectively. In contrast, SEQUEST and Mascot show little advantage in the presence of Byonic: their identified true PSMs are nearly all identified by Byonic (Figure S1). Our results confirm that these five database search algorithms have distinctive advantages in identifying unique PSMs, an observation that aligns well with existing literature [22–26, 39].

In terms of FDR control, four database search algorithms---Byonic, Mascot, SEQUEST, and MS-GF+---demonstrate robust FDR control as they keep the FDPs on the benchmark data under the FDR thresholds *q* ∈ {1%, …,10%}. In contrast, except at small values of *q* such as 1% or 2%, MaxQuant fails the FDR control by a large margin (Figure 1C).

To evaluate the effect of FDR-control procedures on each database search algorithm, we benchmarked two APIR-FDR options, one p-value-based and the other p-value-free, used with each database search algorithm. Specifically, as an exploration, if a database search algorithm uses p-value-based FDR control by default, we used Clipper as an alternative, p-value-free option; otherwise, if the algorithm’s default FDR-control procedure is p-value-free, we used the p-value-based option as an alternative.

On the *Pfu* proteomics standard dataset, we examined the FDPs and power of the five database search algorithms with two APIR-FDR options for a range of FDR thresholds: *q* ∈ {1%, …, 10%}. Our results in Figure 1C show that both p-value-based and p-value-free APIR-FDR options achieve the FDR control and similar power when applied to the outputs of Byonic, Mascot, SEQUEST, and MS-GF+, its default FDR-control procedure, which is the p-value-free FDR-control procedure outlined in the Method section, fails to control the FDR under the target by a large margin. However, the alternative p-value-based FDR-control procedure we applied alleviates the FDR control issue by reducing the FDPs to be closer to the FDR thresholds, with FDPs controlled under *q* when *q* > 5%. Regarding the phenomenon that both the number of true PSMs and the FDP of MaxQuant (with p-value-based FDR control) stay unchanged as the FDR threshold *q* increases from 1% to 10%, we provide a detailed explanation in File S1 and Figure S2.

We also compared the performance of the five database search algorithms with two APIR-FDR options (p-value-based and p-value-free) on the proteomics standard dataset after excluding from each database search algorithm’s output the 1416 shared true PSMs identified by all five algorithms at the FDR threshold *q* = 5%. Our results in Figure S4 show that that default p-value-free FDR-control procedure of MS-GF+ no longer controls the FDR.

Based on the benchmark results above, we chose the p-value-based APIR-FDR option for Maxquant and MS-GF+ because the two algorithms’ default p-value-free FDR-control procedure fail to guarantee the FDR control. For Byonic, Mascot, and SEQUEST, both the p-value-based and p-value-free APIR-FDR options can be used. See Table S1 for details of the APIR-FDR options used with the five database search algorithms in each analysis.

### Set union and intersection operations do not guarantee to control the FDR

In data analysis, there exists a common intuition: if multiple algorithms designed for the same purpose are applied to the same dataset to make discoveries, and all algorithms have their FDRs under *q*, then the intersection of their discoveries (i.e., the discoveries found by all algorithms) should have the FDR under *q* [11]. However, this intuition does not hold in general. The reason is that if all algorithms find different true discoveries, then their common discoveries (i.e., the intersection) could be enriched with false discoveries and thus have the FDR larger than *q*. To demonstrate this, we designed a simulation study called the shared-false-PSMs scenario, where the set intersection operation fails to control the FDR. Although intuition says that the set union operation may not control the FDR, we designed another simulation study called the shared-true-PSMs scenario, where the set union operation fails to control the FDR, for completeness.

Under the shared-true-PSMs scenario, we designed three toy database search algorithms that tend to identify overlapping true PSMs but non-overlapping false PSMs (**Figure 3**A top). In contrast, under the shared-false-PSMs scenario, we designed another three toy database search algorithms that tend to identify overlapping false PSMs but non-overlapping true PSMs (Figure 3A bottom) (see File S1 for the detailed designs of the two scenarios). Under both scenarios, we first applied APIR-FDR to each toy database search algorithm’s output. Then we aggregated identified PSMs from the three algorithms under each scenario using set intersection, set union, or APIR, and we evaluated the FDR of each aggregated PSM set. Figure 3B shows that, while set union fails to control the FDR in the shared-true-PSMs scenario and set intersection fails in the shared-false-PSMs scenario, APIR controls the FDR in both scenarios.

**Figure 3.**
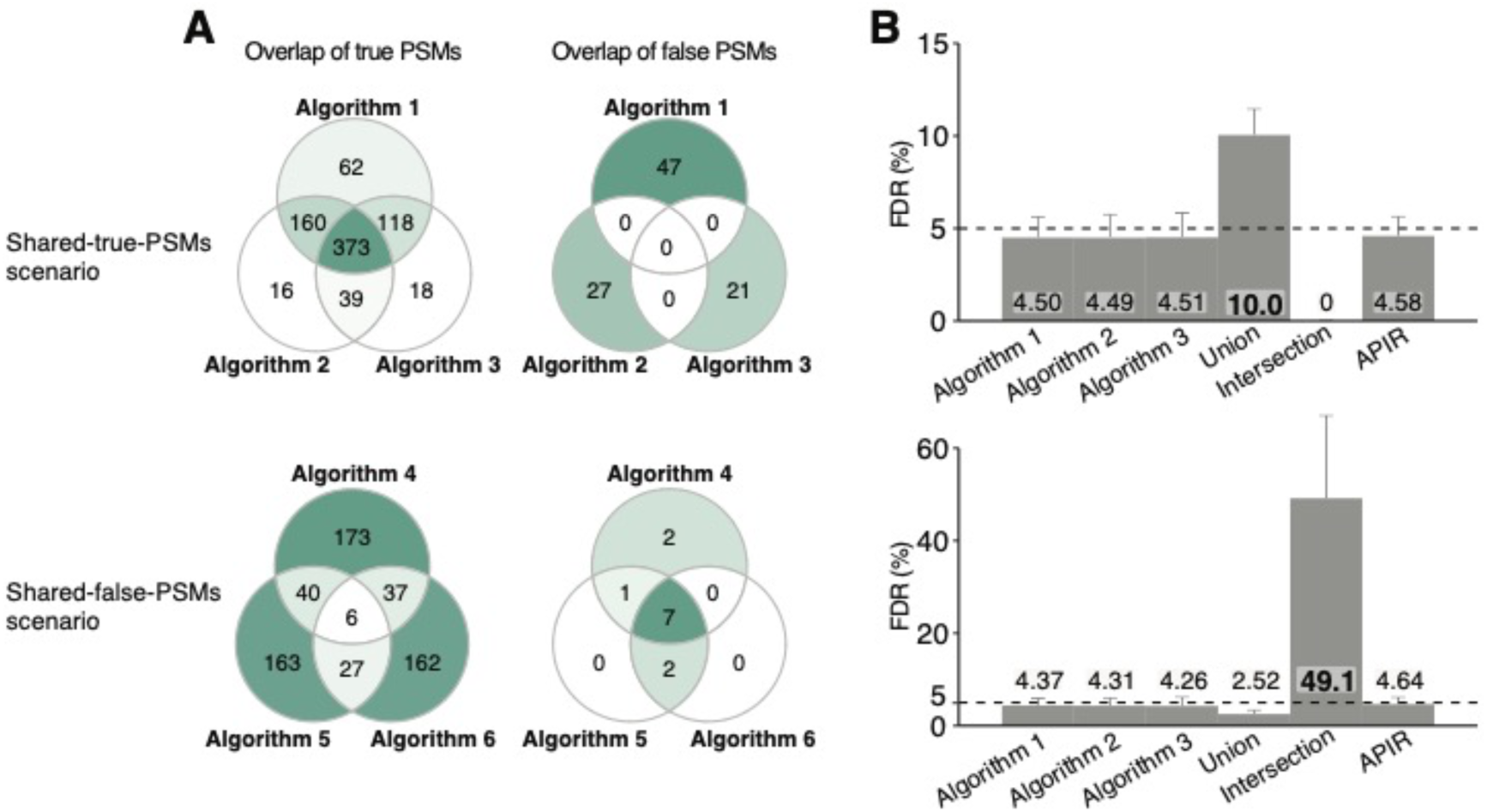
Simulation studies showing that neither intersection nor union of discovery sets (with controlled FDR) controls FDR. FDR control comparison of APIR, intersection, and union for aggregating three toy database search algorithms using simulated data. Two scenarios are considered: the shared-true-PSMs scenario (top) and the shared-false-PSMs scenario (bottom). **A**. Venn diagrams of true PSMs and false PSMs (identified at the FDR threshold *q* = 5%) on one simulation dataset under each simulation scenario. **B**. The FDRs of the three database search algorithms and three aggregation methods: union, intersection, and APIR. Note that the FDR of each database search algorithm or each aggregation method is computed as the average of FDPs on 200 simulated datasets under each scenario.

These two scenarios serve as counterexamples, demonstrating that neither set union nor set intersection can control the FDR of identified target PSMs. In contrast, APIR has the theoretical FDR control.

### APIR has verified FDR control and outpowers Scaffold and ConsensusID

To demonstrate that APIR controls the FDR by aggregating individual search algorithms on the *Pfu* proteomics standard, we benchmarked APIR against two existing aggregation methods, Scaffold and ConsensusID, because they are the only two aggregation methods compatible with the five database search algorithms we used: Byonic, Mascot, SEQUEST, MaxQuant, and MS-GF+. Since database search algorithms are time-consuming to run, we first focused on the 20 combinations consisting of no more than three of the five algorithms, including 10 combinations of any two algorithms and 10 combinations of any three algorithms.

Because of the trade-off between FDR and power, power comparison is valid only when FDR is controlled. Hence, for the three aggregation methods, APIR, Scaffold, and ConsensusID, we compared them in terms of both their FDPs and power on the *Pfu* proteomics standard dataset. Regarding the power increase of each aggregation method over individual database search algorithms, we computed the percentage increases in the aggregated true PSMs, peptides, and proteins by treating as baselines the maximal numbers of true PSMs, peptides, and proteins identified by the five database search algorithms. For example, to aggregate Byonic and MaxQuant, based on our benchmarking results in Figure 1C, we applied (with the default p-value-free FDR-control procedure) and MaxQuant (with p-value-based FDR control) to identify PSMs in Round 1. We would calculate the percentage increase in the identified true PSMs by treating as the baseline the larger of two numbers: the numbers of true PSMs identified by Byonic and MaxQuant.

Our results in **Figure 4** and Figure S5 show that, at both FDR thresholds *q* = 5% and 1%, APIR achieves consistent FDR control and power improvement over individual database search algorithms. In contrast, Scaffold controls the FDR but shows highly inconsistent power improvement, while ConsensusID neither controls the FDR nor has power improvement. Specifically, ConsensusID’s FDPs exceed the FDR threshold *q* = 5% by a large margin: they rise above 15% in 9 out 20 combinations. In summary, only APIR consistently achieves power increase over individual database search algorithms across the 20 algorithm combinations, an advantage that neither Scaffold nor ConsensusID offers.

**Figure 4.**
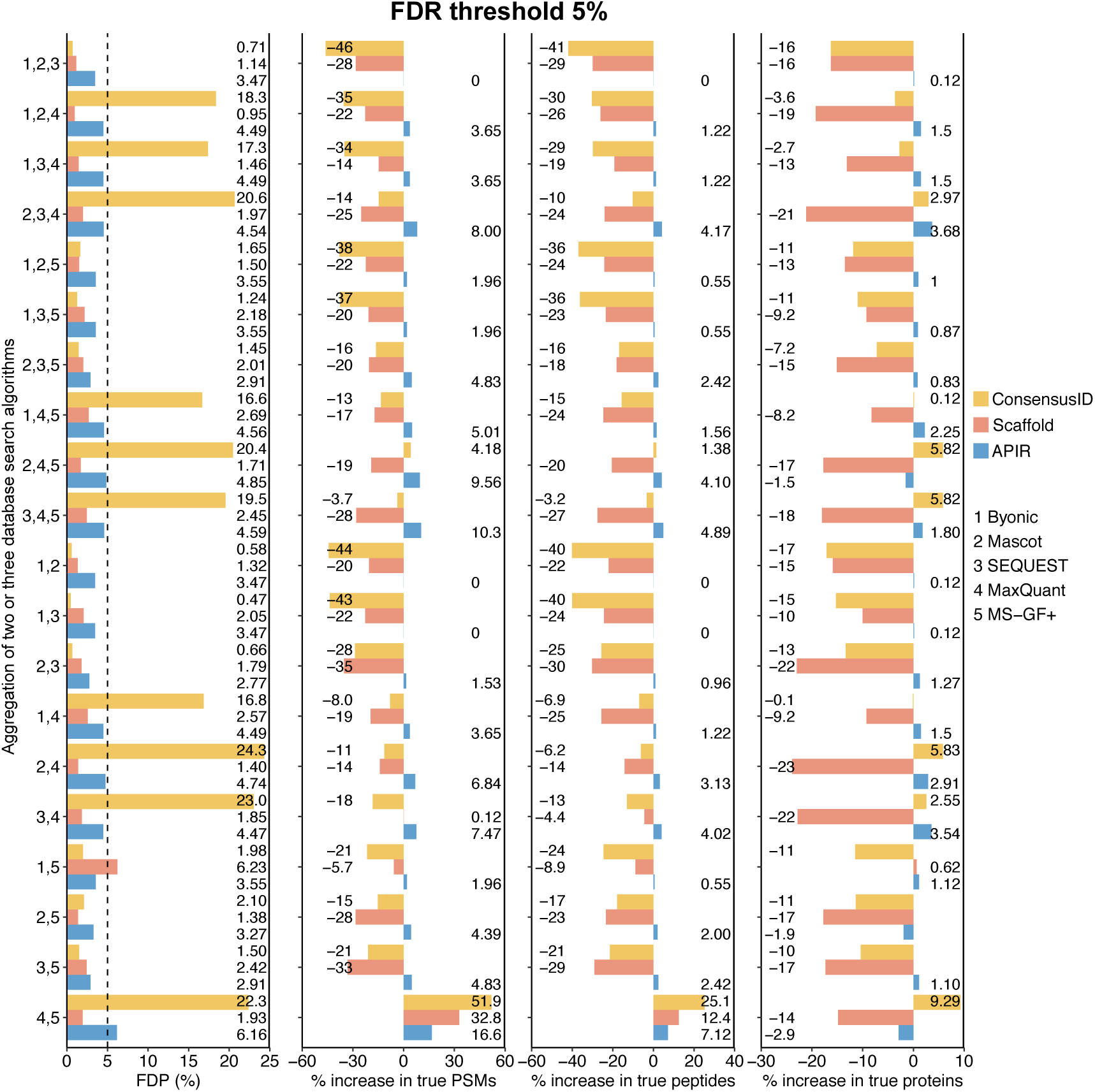
On the proteomics standard dataset, comparison of APIR, Scaffold, and ConsensusID at the FDR threshold *q* = 5%. We set both the peptide threshold and the protein threshold of Scaffold to be 5% FDR. FDPs (first column), the percentage increases in true PSMs (second column), the percentage increases in true peptides (third column), and the percentage increases in true proteins (fourth column) after aggregating two or three database search algorithms out of the five (Byonic, Mascot, SEQUEST, MaxQuant, and MS-GF+). The percentage increase in true PSMs/peptides/proteins is computed by treating as the baseline the maximal number of correctly identified PSMs/peptides/proteins by individual database search algorithms in Round 1 of APIR. Based on the benchmarking results in Figure 1C, in Round 1 of APIR, we applied p-value-free APIR-FDR to Byonic, Mascot, SEQUEST, and MS-GF+, and we applied p-value-based APIR-FDR to MaxQuant. In later rounds of APIR, we used p-value-based APIR-FDR for FDR control.

A technical note is that Scaffold cannot control the FDR of aggregated PSMs; instead, it controls the FDRs of aggregated peptides and proteins, and it requires the FDR thresholds to be input for both. Hence, strictly speaking, Scaffold is not directly comparable with APIR in terms of FDR control because APIR controls the FDR of aggregated PSMs. For a fair comparison, we implemented a variant of Scaffold, which, compared with the default Scaffold, has an advantage in power at the cost of an inflated FDR (see File S1). Our results show that this Scaffold variant demonstrates a slightly inflated FDP in 5 combinations at *q* = 5% (Figure S6A) and 11 combinations at *q* = 1% (Figure S7A). In terms of power, this Scaffold variant still fails to outperform the most powerful individual database search algorithm in 8 combinations at *q* = 5% (Figure S6B) and 10 combinations at *q* = 1% (Figure S7B).

Moreover, we have the results of APIR combining four and five database search algorithms in Figures S8–S9, which again confirm the FDR control and power advantage of APIR. In addition, we examined whether APIR might inflate the peptide-level FDRs by selecting the set of identified PSMs containing the largest number of unique peptides in each round. Figure S10 shows that among the 52 cases (all 26 algorithm combinations × 2 PSM-level FDR thresholds 1% and 5%), APIR either lowers or maintains the maximum peptide-level FDP achieved by an individual search algorithm. In other words, APIR does not inflate the peptide-level FDP.

### APIR empowers peptide identification on AML and TNBC datasets

We next applied APIR with the aforementioned 20 combinations of two and three algorithms to four real datasets: two in-house phospho-proteomics (explained below) AML datasets (“phospho AML-C1” and “phospho AML-C2”) we collected from two cohorts of AML patients (which were not randomly assigned and thus not biological replicates) for studying the properties of LSCs; a published nonphospho-proteomics AML dataset (“nonphospho AML”) that also compares the stem cells with non-stem cells in AML patients [40]; and a published phospho-proteomics TNBC dataset that studies the drug genistein’s effect on breast cancer [41]. Phospho-proteomics is a branch of proteomics; while traditional proteomics aims to capture all peptides in a sample, phospho-proteomics focuses on phosphorylated peptides, also called phosphopeptides, because phosphorylation regulates essentially all cellular processes [42]. See File S1 for the details on how we generated phospho AML-C1 and phospho AML-C2.

On each dataset, we applied APIR at two FDR thresholds *q* = 1% and 5% and examined the percentage increases at four levels: PSMs, peptides, peptides with modifications, and proteins; we calculated the percentage increases in the same way as what we did for the proteomics standard dataset. Our results in **Figure 5** (*q* = 5%) and Figure S11 (*q* = 1%) show that APIR leads to positive percentage increases at two levels (PSMs and peptides) on all four datasets, confirming APIR’s guarantee for identifying more peptides than individual algorithms do. At the peptide-with-modification level, APIR also achieves positive percentage increases across 20 combinations on all four datasets with only one exception: APIR falls short by a negligible 0.1% when aggregating the outputs of Byonic, Mascot, and SEQUEST on the TNBC dataset at *q* = 5%. At the protein level, APIR still manages to outperform individual database search algorithms for all 20 combinations on both phospho-proteomics AML datasets and for more than half of the combinations on the TNBC and nonphospho-proteomics AML datasets. Our results demonstrate that APIR could boost the usage efficiency of shotgun proteomics data.

**Figure 5.**
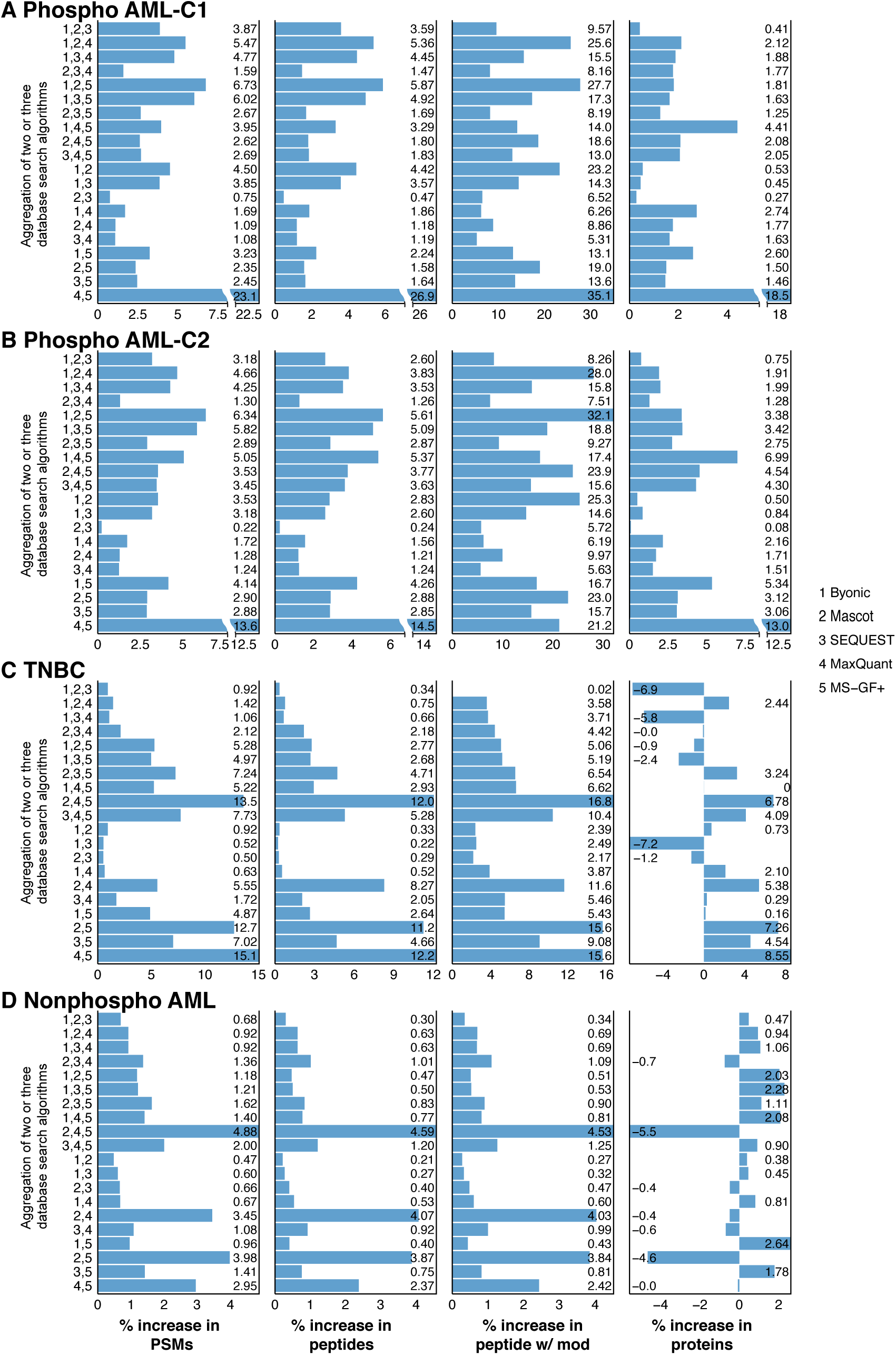
Power improvement of APIR over individual database search algorithms at the FDR threshold *q* = 5%. The percentage increases in PSMs (first column), the percentage increases in peptides (second column), the percentage increases in peptides with modifications (third column), and the percentage increases in true proteins (fourth column) of APIR after aggregating two or three database search algorithms out of the five (Byonic, Mascot, SEQUEST, MaxQuant, and MS-GF+) at the FDR threshold *q* = 5% on (**A**) the phospho AML-C1 dataset, (**B**) the phospho AML-C2 dataset, (**C**) the TNBC dataset, and (**D**) the nonphospho AML dataset. The percentage increase in PSMs/peptides/peptides with modifications/proteins is computed by treating as the baseline the maximal number of PSMs/peptides/peptides and modifications/proteins by an individual database search algorithm in Round 1 of APIR. Phospho AML-C1, phospho-proteomics acute myeloid leukemia-patient cohort 1; phospho AML-C2, phospho-proteomics acute myeloid leukemia-patient cohort 2; TNBC, triple-negative breast cancer; nonphospho AML, nonphospho-proteomics acute myeloid leukemia.

We also applied APIR to combining four and five database search algorithms in Figures S12–S13, which again confirm the power advantage of APIR.

### APIR identifies biologically meaningful proteins from AML and TNBC datasets

Next, we investigated the biological functions of the proteins missed by individual database search algorithms but recovered by APIR from the phospho AML and TNBC datasets. We also performed additional analyses to confirm the existence of these biologically relevant proteins. Specifically, APIR adopts individual search algorithms’ mappings from PSMs to proteins. That is, APIR aggregates PSMs and maps them to proteins based on the PSM-protein mappings output by individual search algorithms. If a PSM is assigned to more than one protein by different search algorithms, APIR outputs a master protein by majority voting. See File S1 for details.

On the phospho AML-C1 and AML-C2 datasets, which contain patient samples with enriched or depleted LSCs, APIR identifies biologically relevant proteins that were missed by individual database search algorithms. Specifically, on phospho AML-C1, APIR identifies from the 20 combinations (of two and three algorithms) 80 additional proteins (the union of the additional proteins APIR identified from the combinations) at the FDR threshold *q* = 1% and 121 additional proteins at the FDR threshold *q* = 5%. These two sets of additional proteins recovered by APIR include some well-known proteins, such as transcription intermediary factor 1-alpha (TIF1α), phosphatidylinositol 4,5-bisphosphate 5-phosphatase A (PIB5PA), homeobox protein Hox-B5 (HOXB5), small ubiquitin-related modifier 2 (SUMO-2), transcription factor jun-D (JUND), glypican-2 (GPC2), dnaJ homolog subfamily C member 21 (DNAJC21), mRNA decay activator protein ZFP36L2. Here we summarize the tumor-related functions of these well-known proteins. High levels of TIF1α are associated with oncogenesis and disease progression in a variety of cancer lineages such as AML [43–49]. PIB5PA has a tumor-suppressive role in human melanoma [50]. Its high expression is correlated with limited tumor progression and better prognosis in breast cancer patients [51]. HOXB5 is among the most affected transcription factors by the genetic mutations that initiate AML [52–54]. SUMO-2 plays a key role in regulating CBX2, which is overexpressed in several human tumors (e.g., leukemia) and whose expression is correlated with lower overall survival [55]. JUND plays a central role in the oncogenic process leading to adult T-cell leukemia [56]. GPC2 is an oncoprotein and a candidate immunotherapeutic target in high-risk neuroblastoma [57]. DNAJC21 mutations are linked to cancer-prone bone marrow failure syndrome [58]. ZFP36L2 induces AML cell apoptosis and inhibits cell proliferation [59]; its mutation is associated with the pathogenesis of acute leukemia [60]. Moreover, on phospho AML-C2, APIR identifies 62 additional proteins at FDR 1% and 19 additional proteins at FDR 5%, including JUND and myeloperoxidase (MPO). MPO is expressed in hematopoietic progenitor cells in prenatal bone marrow, which are considered initial targets for the development of leukemia [61–63].

On the TNBC dataset, APIR identifies 92 additional proteins missed by individual database search algorithms at the FDR threshold *q* = 1% and 69 additional proteins at *q* = 5%. In particular, at *q* = 1%, APIR uniquely identifies breast cancer type 2 susceptibility protein (BRCA2) and Fanconi anemia complementation group E (FANCE). BRCA2 is a well-known breast cancer susceptibility gene; an inherited genetic mutation inactivating the BRCA2 gene is found in TNBC patients [64–69]. The FANC-BRCA pathway, including FANCE and BRCA2, is known for its roles in DNA damage response. Inactivation of the FANC–BRCA pathway is identified in ovarian cancer cell lines and sporadic primary tumor tissues [70, 71]. Additionally, at both *q* = 1% and 5%, APIR identifies JUND and roundabout guidance receptor 4 (ROBO4); the latter regulates tumor growth and metastasis in multiple types of cancer, including breast cancer [72–75]. We summarize the biological relevance of these proteins in **Table 1**.

**Table 1:**
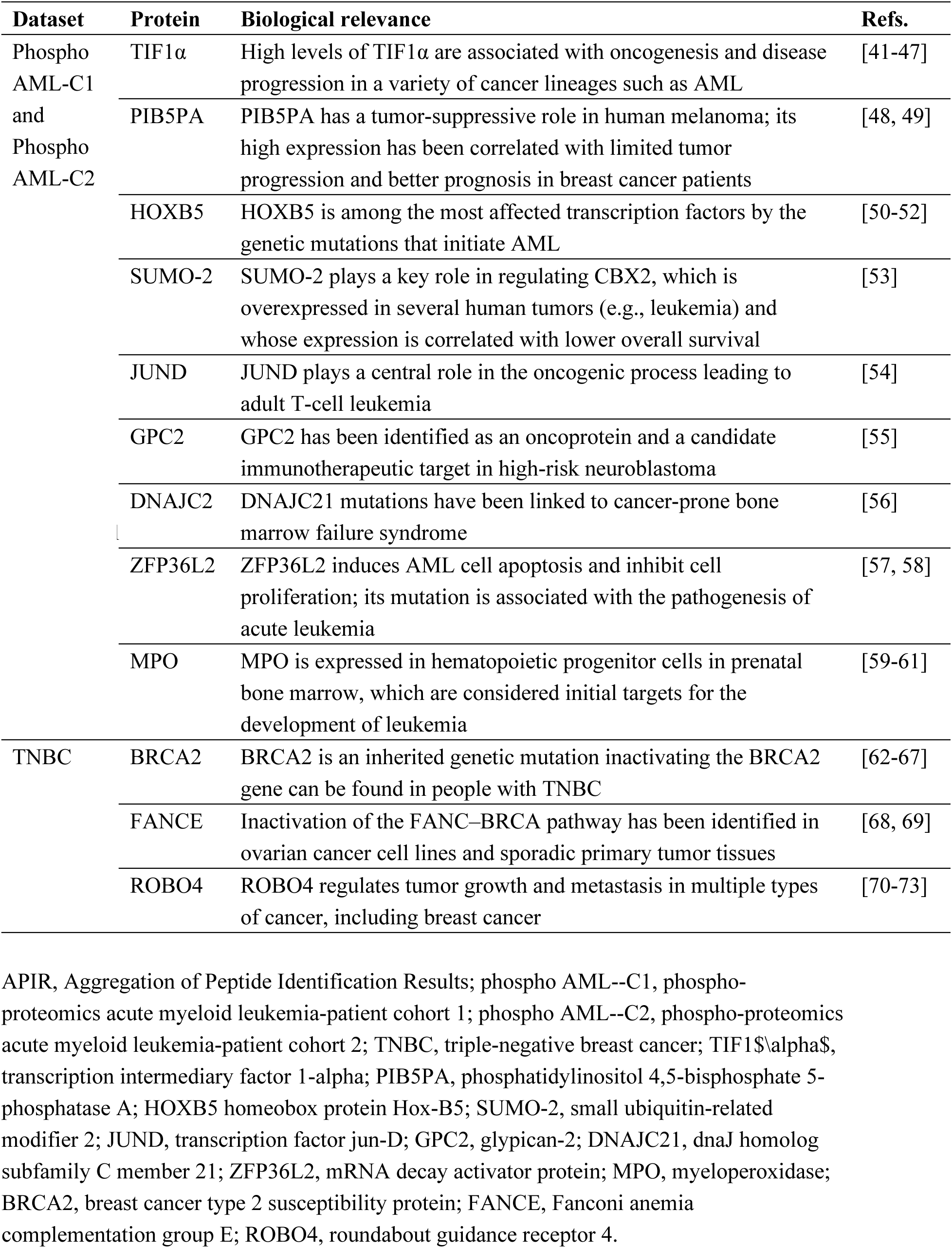
A summary of biologically relevant proteins missed by individual database search algorithms but recovered by APIR from the phospho AML--C1 and AML--C2 and TNBC datasets.

To further evaluate the existence of the aforementioned known proteins, we performed two analyses. First, we examined the tandem MS spectra of the PSMs corresponding to these proteins identified from the phospho AML datasets. Our results in Table S2 and File S2 show that the PSMs rescued by APIR are likely true positives. The rescued PSMs fell broadly into three categories: 1) high-likelihood identifications with both accurate precursor mass and numerous fragment ions (40%), 2) identifications based on accurate precursor mass and few (30%) or no fragment ions (10%), and 3) chimeric spectra (20%). Second, we examined the PSMs corresponding to these proteins identified from the phospho AML datasets and the TNBC dataset (Tables S4–S6), and we found that these proteins all correspond to at least one target PSM with a high matching score (from at least one database search algorithm). These results, combined with the constituent nature and biological relevance of these proteins (Table 1), suggest the likely existence of these proteins and demonstrate APIR’s potential in identifying novel disease-related proteins.

### APIR empowers the identification of differentially expressed peptides

An important use of proteomics data is the differential expression (DE) analysis, which aims to identify proteins whose expression levels change between two conditions. Protein is the ideal unit of measurement; however, due to the difficulties in quantifying protein levels from tandem MS data, an alternative approach has been proposed and used, which first identifies differentially expressed peptides and then investigates their corresponding proteins along with modifications. Because it is less error-prone to quantify peptides than proteins, doing so would dramatically reduce errors in the DE analysis.

We compared APIR with MaxQuant and MS-GF+ by performing DE analysis on the phospho AML-C1 dataset. We focused on this dataset instead of the TNBC dataset or the nonphospho AML dataset because the phospho AML datasets were generated for our in-house study and thus may yield new discoveries. This analysis is conducted to demonstrate that APIR could improve the identification power by aggregating dissimilar algorithms. Since MaxQuant and MS-GF+ identify drastically different PSMs on our real datasets (Figure S14) and are widely-used, open-source tools, we selected them as two example algorithms.

The phospho AML-C1 dataset contains six bone marrow samples: three enriched with LSCs, two depleted of LSCs, and one control. To simplify our DE analysis, we selected two pairs of enriched and depleted samples. Specifically, we first applied APIR to aggregate the outputs of MaxQuant and MS-GF+ on the phospho AML-C1 dataset using all six samples. Then we applied DESeq2 to identify DE peptides from the aggregated peptides of APIR, MaxQuant, and MS-GF+ using the four selected samples.

Our results in **Figure 6** show that at the FDR threshold *q* = 5%, we identified 318 DE peptides from 224 proteins based on APIR, 251 DE peptides from 180 proteins based on MaxQuant, and 242 DE peptides from 190 proteins based on MS-GF+, respectively. In particular, APIR identified 6 leukemia-related proteins: the promyelocytic leukemia zinc finger (PLZF), serine/threonine-protein kinase B-raf (B-raf), signal transducer and activator of transcription 5B (STAT5B), promyelocytic leukemia protein (PML), cyclin-dependent kinase inhibitor 1B (CDKN1B), and retinoblastoma-associated protein (RB1), all of which belong to the AML KEGG pathway or the chronic myeloid leukemia KEGG pathway [76–78]. In particular, PLZF and CDKN1B were uniquely identified from the APIR aggregated results but not by either MaxQuant or MS-GF+.

**Figure 6.**
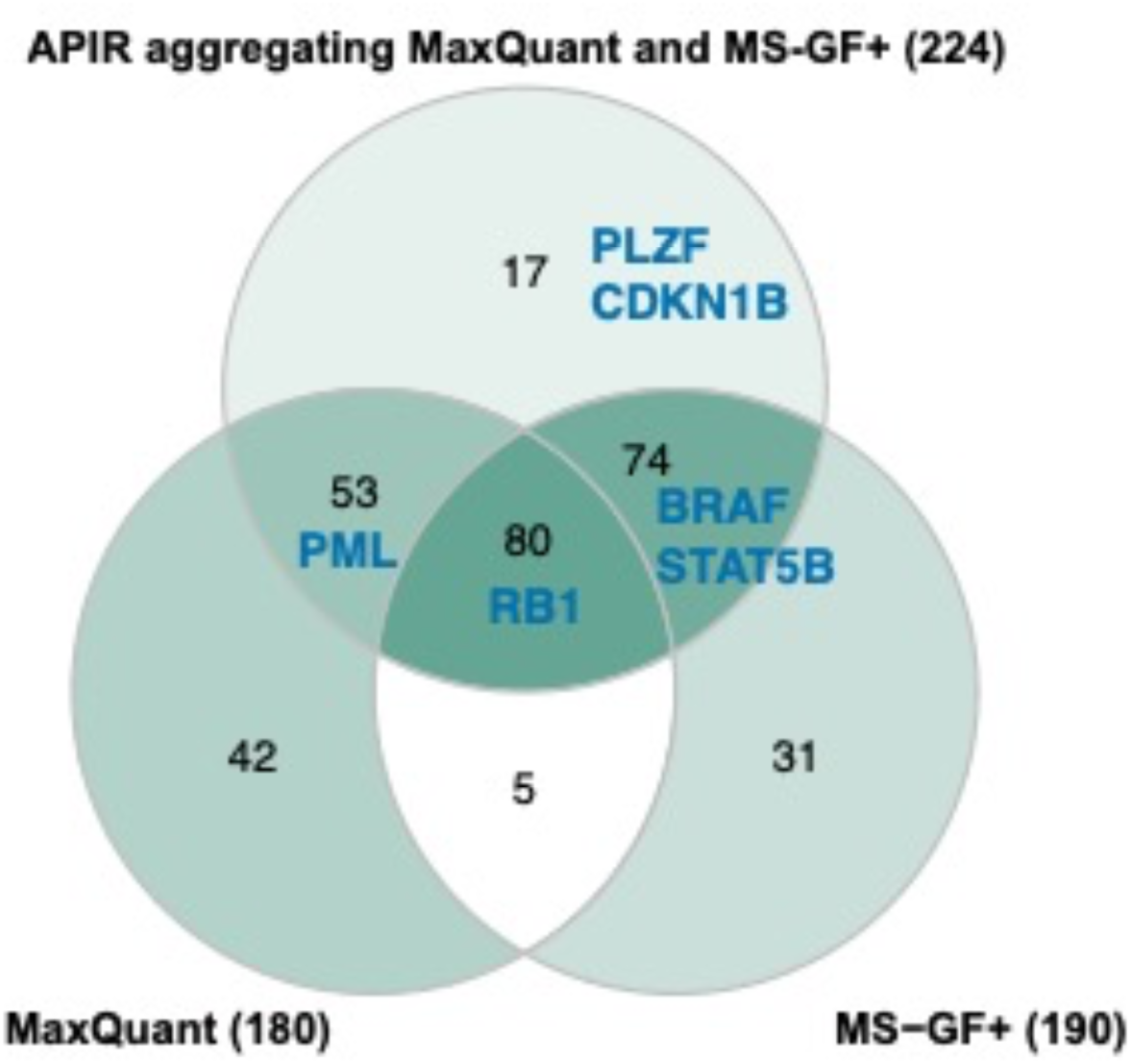
Comparison of APIR with MaxQuant and MS-GF+ by DE analysis on the phospho AML-C1 dataset. Venn diagrams of DE proteins based on the identified peptides by APIR aggregating MaxQuant and MS-GF+, MaxQuant, and MS-GF+. Six leukemia-related proteins were found as DE proteins based on APIR: PLZF, B-raf, STAT5B, PML, CDKN1B, and RB1. Notably, this dataset contains six bone marrow samples from two patients: P5337 and P5340. From P5337, one LSC-enriched sample and one LSC-depleted sample were taken. From P5340, two LSC-enriched samples and one LSC-depleted sample were taken. In our DE analysis, we compare two LSC-enriched samples (one per patient) against two LSC-depleted samples (one per patient). DE, differentially expressed; LSC, leukemia stem cells.

We next investigated the phosphorylation of the identified DE peptides of PLZF or CDKN1B. With regard to PLZF, APIR identified phosphorylation at Threonine 282, which is known to activate cyclin-A2 [79], a core cell cycle regulator of which the deregulation seems to be closely related to chromosomal instability and tumor proliferation [80–82]. As for CDKN1B, APIR identified phosphorylation at Serine 140. Previous studies have revealed that ATM phosphorylation of CDKN1B at Serine 140 is important for stabilization and enforcement of the CDKN1B-mediated G1 checkpoint in response to DNA damage [83]. A recent study shows that inability to phosphorylate CDKN1B at Serine 140 is associated with enhanced cellular proliferation and colony formation [84]. Our results, summarized in **Table 2**, demonstrate that APIR can assist in discovering interesting proteins and relevant post-translational modifications.

**Table 2:** A summary of biologically relevant phosphorylation sites in the DE peptides identified by DESeq2 from the aggregated peptides by APIR from the outputs of MaxQuant and MS-GF+ on the phospho AML--C1 dataset.

| **Protein** | **Phosphorylation site** | **Biological relevance** | **Refs.** |
| --- | --- | --- | --- |
| PLZF | Thr 282 | Phosphorylation at Threonine 282 activates cyclin-A2, a core cell cycle regulator of which the deregulation seems to be closely related to chromosomal instability and tumor proliferation | [77-80] |
| CDKN1B | Ser 140 | Phosphorylation of CDKN1B at Serine 140 is important for stabilization and enforcement of the CDKN1B-mediated G1 checkpoint in response to DNA damage; inability to phosphorylate CDKN1B at Serine 140 is associated with enhanced cellular proliferation and colony | [81, 82] |

DE, differentially expressed; PLZF, the promyelocytic leukemia zinc finger; CDKN1B, cyclin-dependent kinase inhibitor 1B.

## Discussion

We developed a statistical framework APIR to combine the power of distinct database search algorithms by aggregating their identified PSMs from shotgun proteomics data with FDR control. The core component of APIR is APIR-FDR, an FDR-control method that re-identifies PSMs from a single database search algorithm’s output without restrictive distribution assumptions. APIR offers a great advantage of flexibility: APIR is compatible with any database search algorithms. The reason lies in that APIR is a sequential approach based on a mathematical fact: given multiple disjoint sets of discoveries, each with the FDP smaller than or equal to *q*, their union also has the FDP smaller than or equal to *q*. This sequential approach not only allows APIR to circumvent the need to impose restrictive distribution assumptions on each database search algorithm’s output, but also ensures that APIR would identify at least as many, if not more, unique peptides as a single database search algorithm does.

By assessing APIR on the first publicly available complex proteomics standard dataset we generated, we verified that APIR consistently improves the power of peptide identification with the FDR controlled on the identified PSMs. Our extensive studies on AML and TNBC data suggest that APIR can discover additional disease-relevant peptides and proteins that are otherwise missed by individual database search algorithms.

We note that [29] developed a multi-stage method to combine PSMs identified by multiple database search algorithms, a seemingly similar framework. However, three major differences exist between APIR and the multi-stage method in [29]. First, APIR is an open-source and platform-agnostic framework that is universally compatible with all database search algorithms. In contrast, the multi-stage method is restricted to three database search algorithms: X!Tandem [85], InsPecT [86], and SpectraST [87]. Second, APIR adopts a data-driven approach to determine the combination order of database search algorithms (Figure 2). In contrast, the multi-stage method pre-determines the combination order of its three database search algorithms based on domain knowledge, making its generalization to other database search algorithms non-trivial. In particular, the last paragraph of [29] says, *“We note, however, that routine application of iterative strategies such as the one utilized in this work, especially in a high throughput environment, will require further substantial work on the development of statistical FDR estimation methods applicable to a wide range of peptide identification approaches, including subset database searching, blind PTM analysis, and genomic searches.”* Hence, APIR makes contribution to the future work mentioned in [29].

The current implementation of APIR controls the FDR at the PSM level. However, in shotgun proteomics experiments, PSMs serve merely as an intermediate to identify peptides and then proteins, the real molecules of biological interest; thus, an ideal FDR control should occur at the protein level. A fact is that FDR control at the PSM level does not entail FDR control at the protein level because multiple PSMs may correspond to the same peptide sequence, and multiple peptides may correspond to the same protein. To realize the FDR control on the identified proteins, APIR-FDR needs to be carefully modified. A possible modification would be to construct a matching score for each protein from the matching scores of the PSMs that correspond to this protein’s peptides. Future studies are needed to explore possible ways of constructing proteins’ matching scores. Once we modify APIR-FDR to control the FDR at the protein level, the current sequential approach of APIR still applies: applying the modified APIR-FDR to sequentially identify disjoint sets of proteins from individual database search algorithms’ outputs; outputting the union of these disjoint sets as discoveries.

Notably, APIR adopts a statistical inference framework as opposed to a machine learning prediction framework for PSM aggregation. Hence, APIR is unlike existing machine learning methods (such as PepArML [11]), which could be categorized into two types. Methods of the first type require an external benchmark proteomics dataset, which contains known true PSMs and false PSMs, as the training data to train a classifier. Then they apply the trained classifier to a new proteomics dataset to predict whether a target PSM is true or false. Their underlying assumption is that the classifier trained on the benchmark dataset is generalizable to the new dataset. However, when this generalizability does not hold (a likely scenario given the vast diversity of biological samples), their predicted target PSMs would become questionable. Methods of the second type do not rely on an external benchmark dataset but have to label a subset of target PSMs as positive or negative for training a classifier. This labelling step requires multiple arbitrary thresholds, which would affect the classifier’s prediction accuracy. In contrast, APIR requires no external training data or arbitrary labelling.

Although the applications in this work are based on tandem MS data collected by data-dependent acquisition (DDA), APIR is also applicable to tandem MS data collected by data-independent acquisition (DIA), as long as the database search algorithms use the target-decoy search strategy. Moreover, although APIR is designed for proteomics data, its framework is general and extendable to aggregating discoveries in other popular high-throughput biomedical data analyses, including peak calling from ChIP-seq data, differential gene expression analysis from bulk or single-cell RNA sequencing data, and differentially interacting chromatin region identification from Hi-C data [32]. For example, an extended APIR may aggregate discoveries made by popular differential gene expression analysis methods, such as DESeq2 [88], edgeR [89], and limma [90] to strengthen FDR control [91] and meanwhile increase the power.

## Data availability

The *Pfu* mass spectrometry dataset is available at the PRoteomics IDEntifications Database (PRIDE) [92] with the dataset identifier PXD028558.

## Code availability

The APIR R package is available at https://github.com/yiling0210/APIR or https://ngdc.cncb.ac.cn/biocode/tools/BT007298. The code and preprocessed data for reproducing the figures are available at https://doi.org/10.5281/zenodo.5202768.

## CRediT author statement

**Yiling Elaine Chen**: Methodology, Software, Formal analysis, Writing – Original Draft, Writing – Review & Editing, Visualization. **Kyla Woyshner**: Software, Formal analysis, Writing – Original Draft. **MeiLu McDermott**: Software, Formal analysis, Writing – Original Draft. **Antigoni Manousopoulou**: Investigation, Writing – Original Draft. **Scott B. Ficarro**: Investigation, Validation. **Jarrod A. Marto**: Investigation. **Xinzhou Ge**: Writing – Review & Editing. **Leo David Wang**: Conceptualization, Resources. **Jingyi Jessica Li**: Conceptualization, Methodology, Writing – Review & Editing.

## Acknowledgments

The authors appreciate the comments and feedback from all members of the Junction of Statistics and Biology at the University of California, Los Angeles, USA (http://jsb.ucla.edu). We also thank the editor and the anonymous reviewers for their insightful comments and suggestions.

This work was supported by the following grants: the National Cancer Institute (a part of the National Institutes of Health in the USA under Grant No. T32LM012424) to YEC; the National Cancer Institute (under Grant No. K08CA201591), Margaret Early Memorial Research Trust in the USA, and Pediatric Cancer Research Foundation in the USA to LDW; the National Cancer Institute (under Cancer Center Support Grant and grant No. P30CA033572) to the mass spectrometry facility at the City of Hope; the National Institute of General Medical Sciences (a part of the National Institutes of Health, USA, under Grant No. R01GM120507 and R35GM140888), the National Science Foundation in the USA (under Grant No. DBI-1846216 and DMS-2113754), Johnson & Johnson WiSTEM2D Award in the USA, Sloan Research Fellowship in the USA, and UCLA David Geffen School of Medicine W.M. Keck Foundation Junior Faculty Award in the USA, to JJL.

## ORCID

0000-0001-7259-2787 (Yiling Elaine Chen)

0000-0002-1005-9896 (Kyla Woyshner)

0000-0002-0866-0738 (MeiLu McDermott)

0000-0001-5028-1865 (Antigoni Manousopoulou)

0000-0002-1521-7996 (Scott B. Ficarro)

0000-0003-2086-1134 (Jarrod A. Marto)

0000-0002-8229-5716 (Xinzhou Ge)

0000-0002-2945-9005 (Leo David Wang)

0000-0002-9288-5648 (Jingyi Jessica Li)

## Supplementary Figures

**Figure S1.**
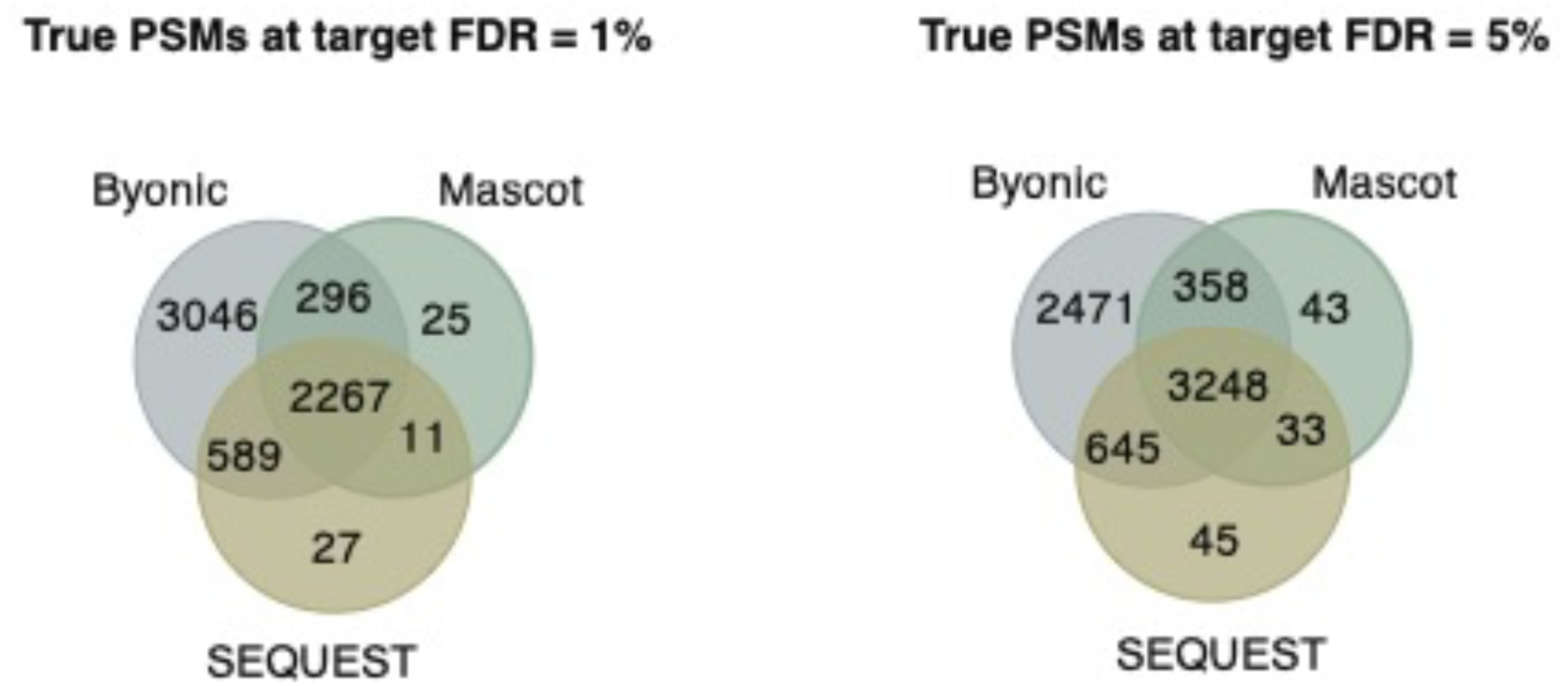
Overlaps of true PSMs identified by Byonic, Mascot, and SEQUEST. Venn diagrams of true PSMs identified by the three database search algorithms from Proteome Discoverer™ Software—Byonic, Mascot, and SEQUEST—under the FDR threshold *q* = 1% (left) or *q* = 5% (right) on the proteomics standard dataset. The true PSMs identified by Byonic nearly cover the true PSMs identified by Mascot or SEQUEST.

**Figure S2.**
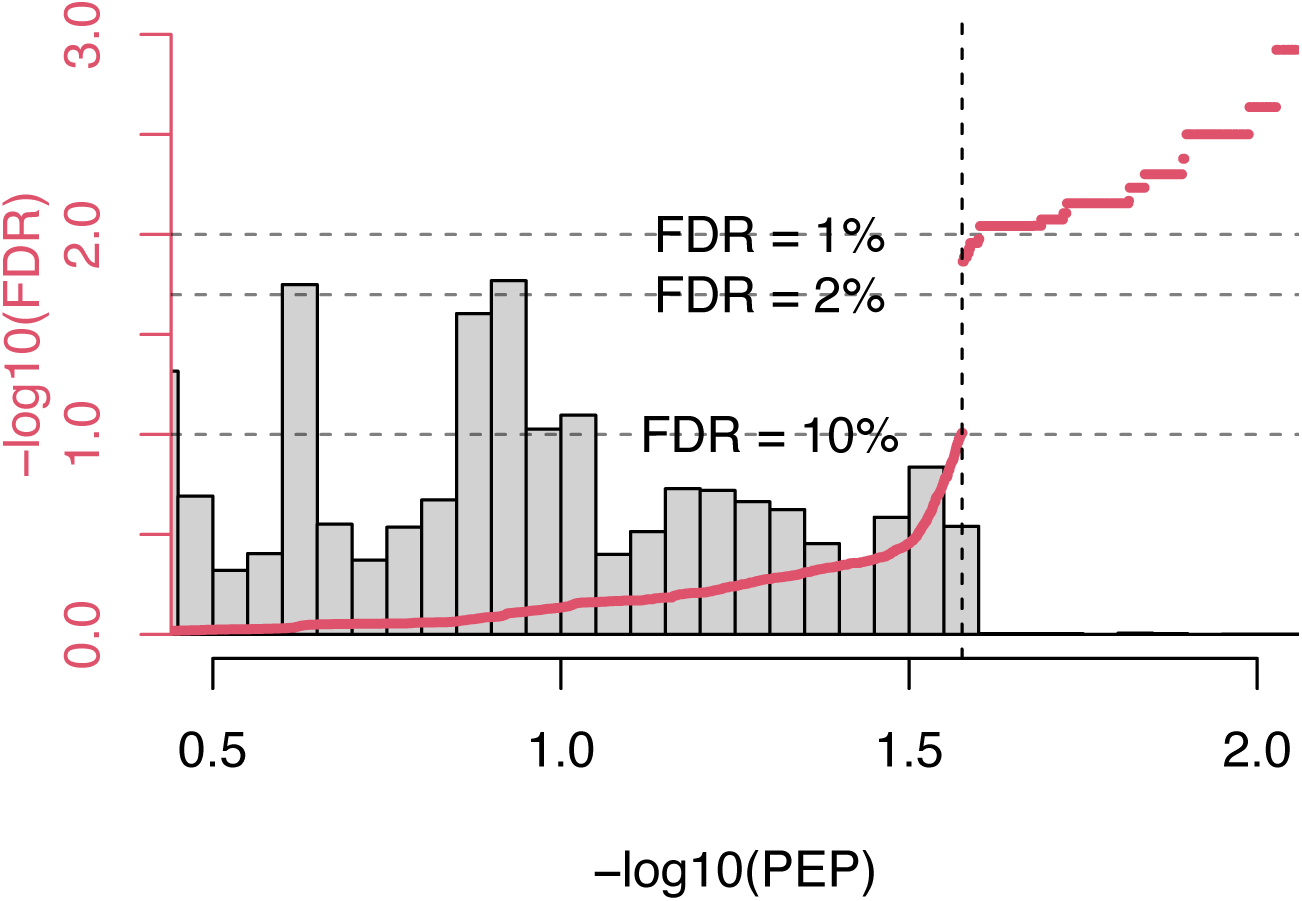
The change of − log_10_ −transformed FDR estimate by APIR (y-axis) with respect to the matching scores of target PSMs by MaxQuant (x-axis) on the proteomics standard dataset. We used − log_10_ −transformed PEP output by MaxQuant as the matching scores. The underlying histogram represents the distribution of matching scores of the decoy PSMs by MaxQuant.

**Figure S3.**
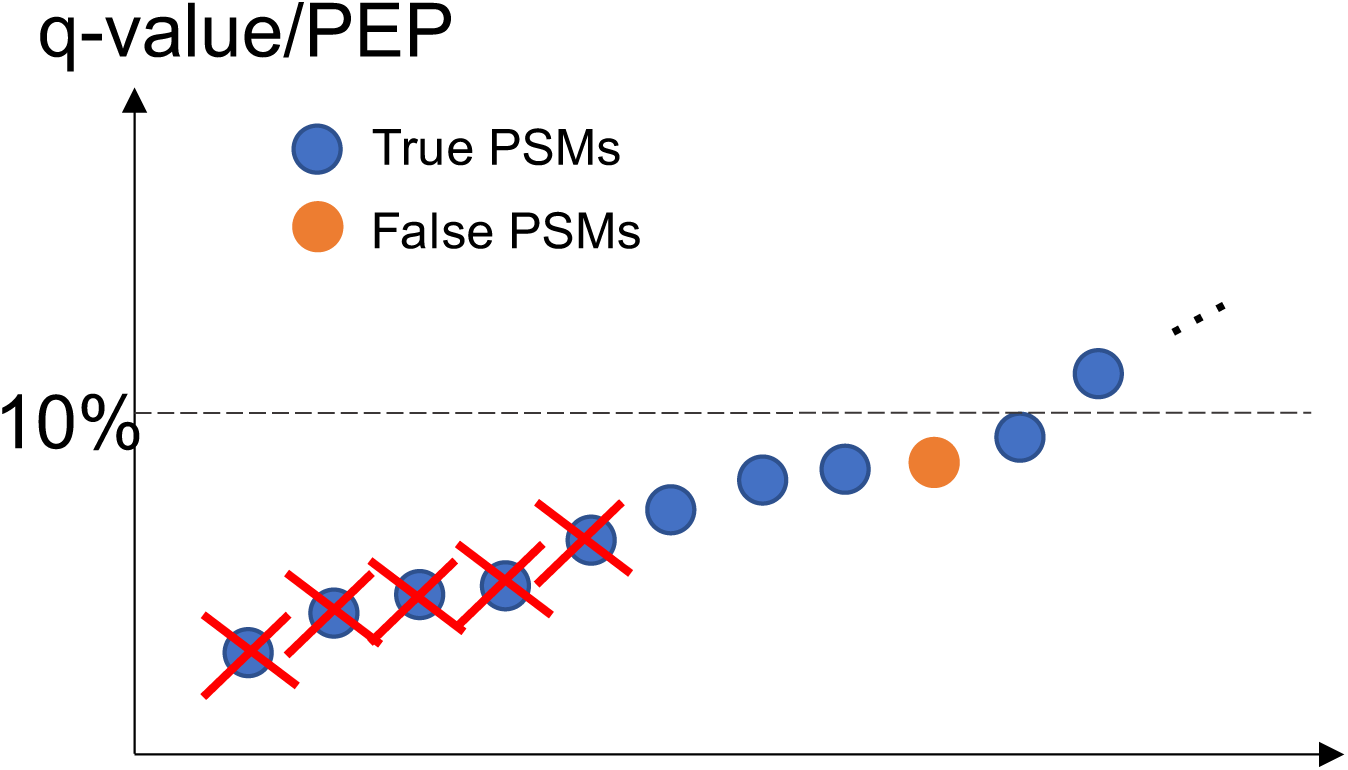
A counter-example explaining why p-value-free FDR control procedure no longer guarantees to control the FDR after a subset of PSMs is removed. Suppose that before removing PSMs, at the FDR threshold 10%, a database search algorithm reports 10 PSMs, whose estimated FDRs (q-values or PEPs) are under 10%, as discoveries. Among these 10 PSMs, 1 is false (in orange), and 9 are true (in blue), so the actual FDP is 10%. In contrast, suppose that we first remove the 5 PSMs with the smallest q-values or PEPs (crossed out) and next threshold the remaining PSMs at the q-value or PEP threshold 10%. Then the actual FDP becomes 1/5 = 20%.

**Figure S4.**
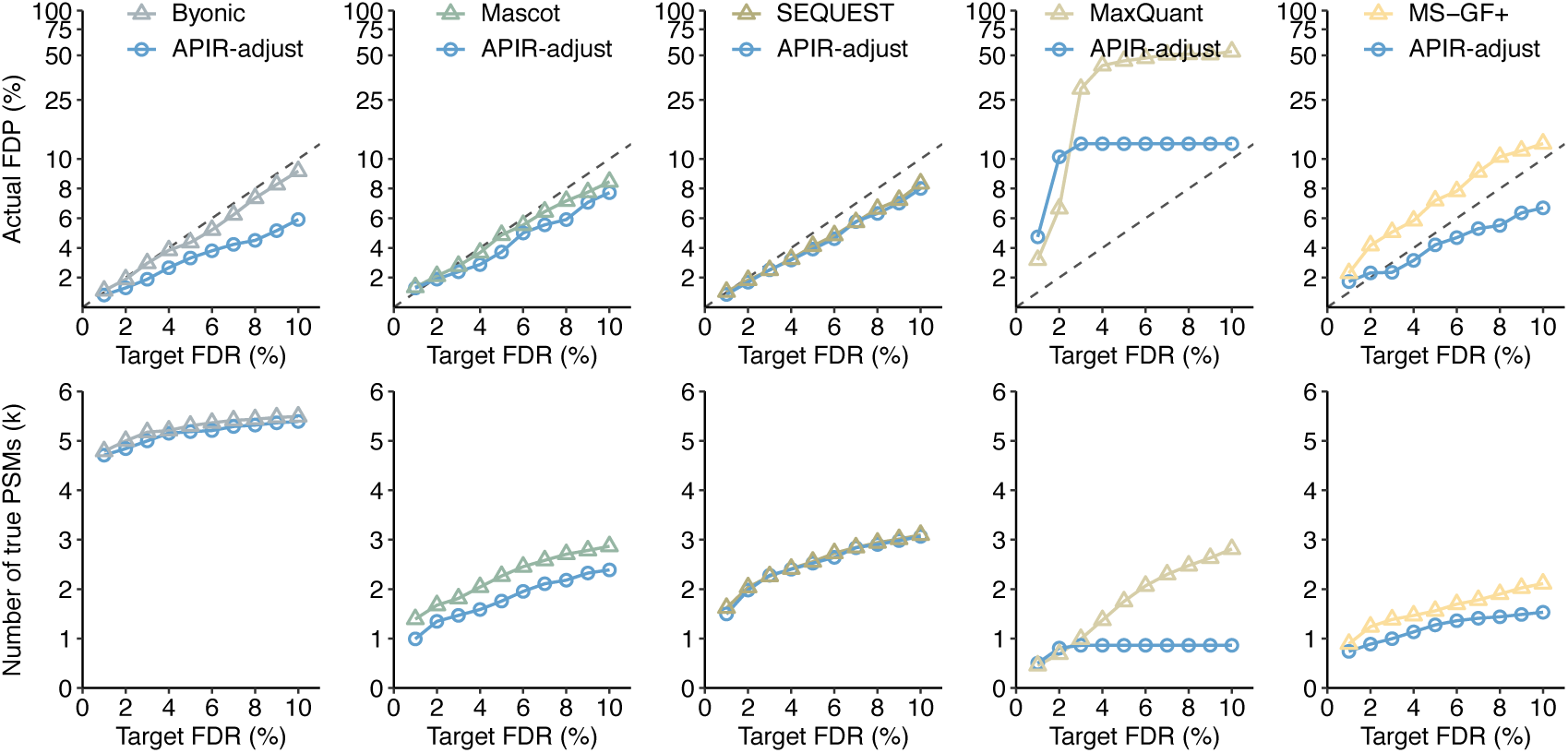
Comparing APIR-FDR options on incomplete output from database search algorithms. At the FDR threshold *q* ∈ {1%, …, 10%}, FDPs and power of each of the five database search algorithms when the 1416 target PSMs (identified by all five database search algorithms at the FDR threshold *q* = 5%) are removed from the output of database search algorithms.

**Figure S5.**
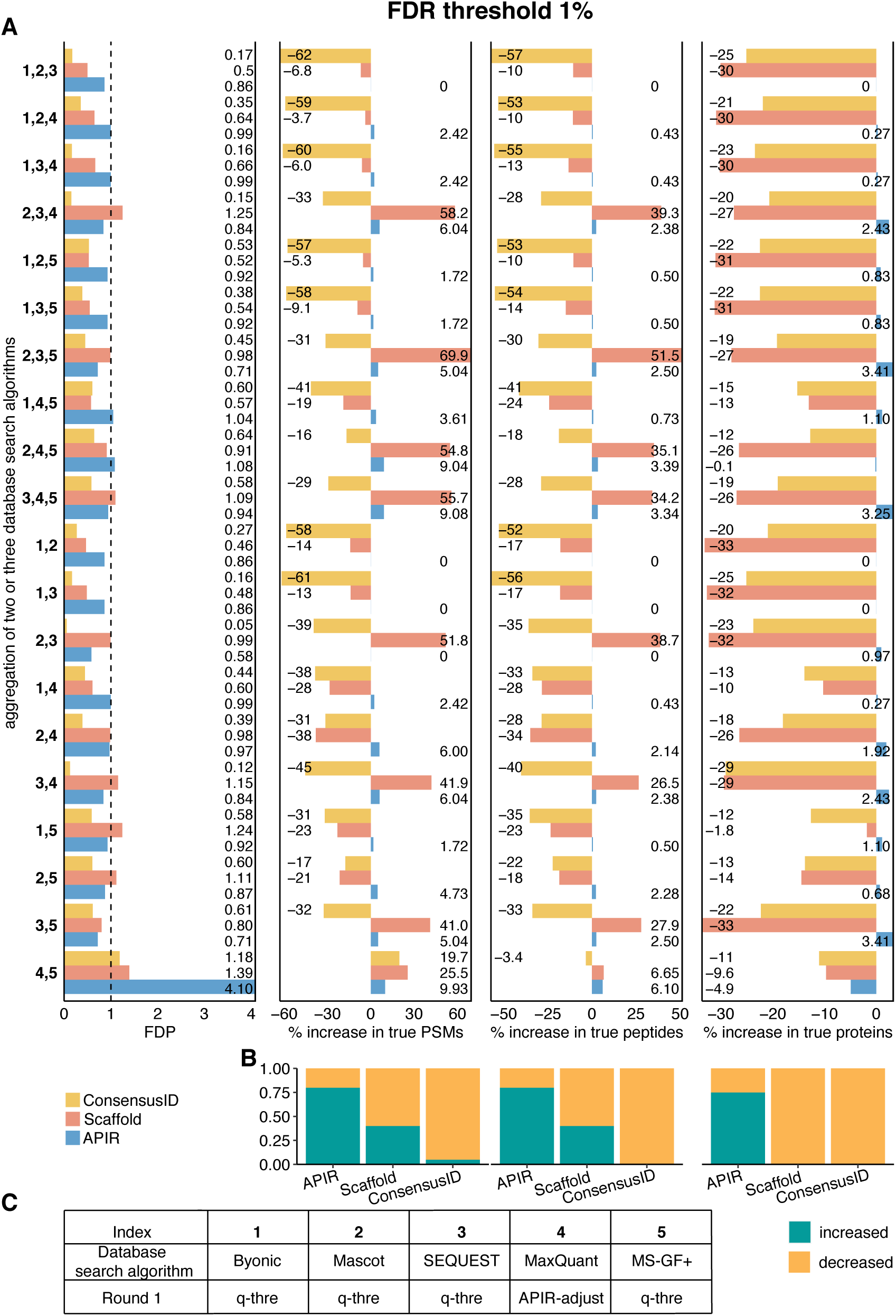
On the proteomics standard dataset, comparison of APIR, Scaffold, and ConsensusID at the FDR threshold *q* = 1% in terms of FDR control and power. We set both the peptide threshold and the protein threshold of Scaffold to be 1% FDR. **A.** FDPs (first column), the percentage increases in true PSMs (second column), the percentage increases in true peptides (third column), and the percentage increases in true proteins (fourth column) after aggregating two or three database search algorithms out of the five (Byonic, Mascot, SEQUEST, MaxQuant, and MS-GF+). The percentage increase in true PSMs/peptides/proteins is computed by treating as the baseline the maximal number of correctly identified PSMs/peptides/proteins by individual database search algorithms in Round 1 of APIR. **B**. Proportions of combinations that show a non-negative percentage increase (green bars) in true PSMs (first column), true peptides (second column), and true proteins (third column). **C**. The indices of database search algorithms in (A) and the implementation of APIR in Round 1. Based on the benchmarking results in Figure 1C, in Round 1 of APIR, we applied p-value-free APIR-FDR to Byonic, Mascot, SEQUEST, and MS-GF+, and we applied p-value-based APIR-FDR to MaxQuant. In later rounds of APIR, we used p-value-based APIR-FDR for FDR control.

**Figure S6.**
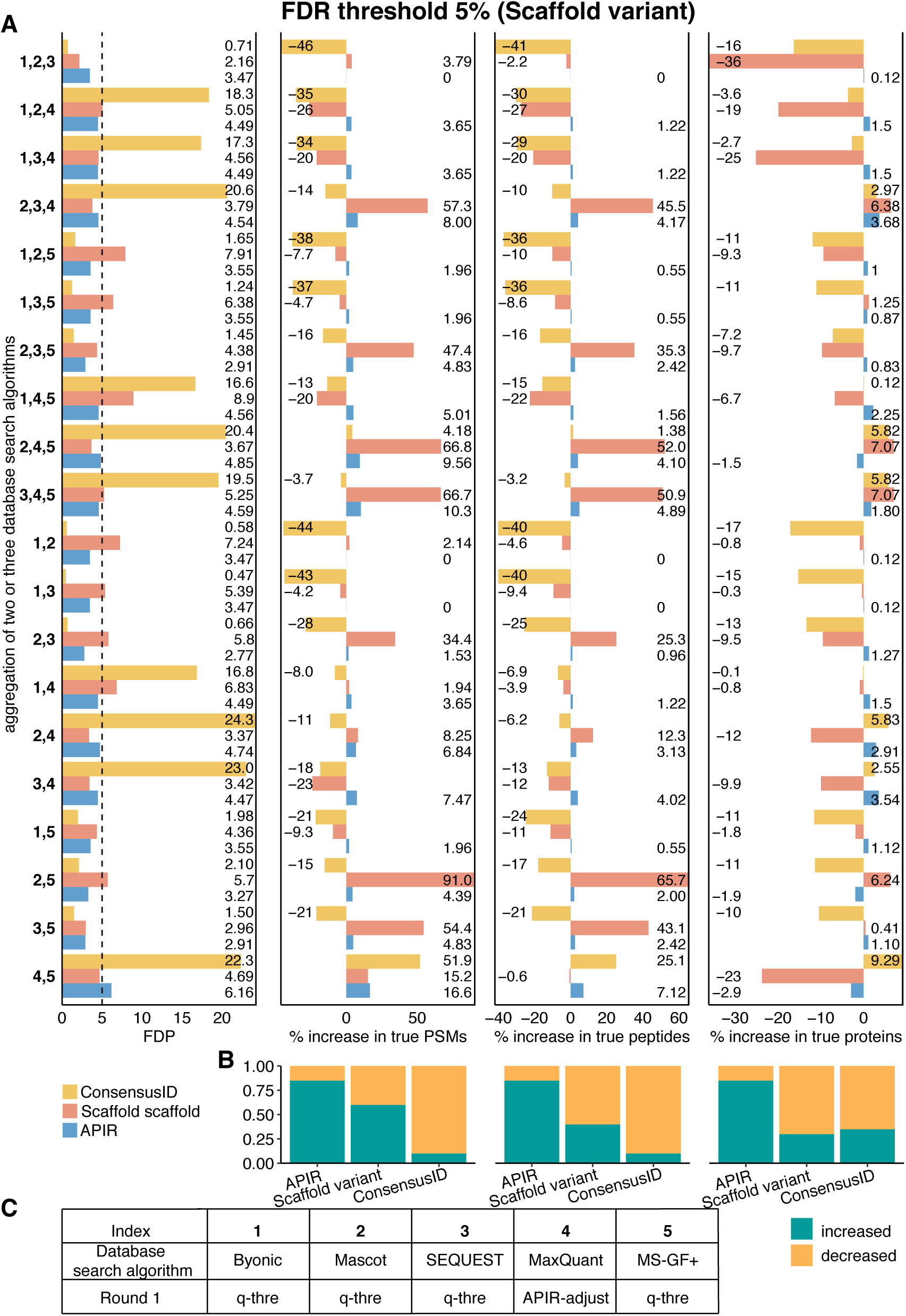
On the proteomics standard dataset, comparison of APIR, Scaffold variant, and ConsensusID at the FDR threshold *q* = 5% in terms of FDR control and power. We set Scaffold’s peptide threshold to be 5% FDR and varied its protein threshold to find the maximal number of identified peptides. **A**. FDPs (first column), the percentage increases in true PSMs (second column), the percentage increases in true peptides (third column), and the percentage increases in true proteins (fourth column) after aggregating two or three database search algorithms out of the five (Byonic, Mascot, SEQUEST, MaxQuant, and MS-GF+). The percentage increase in true PSMs/peptides/proteins is computed by treating as the baseline the maximal number of correctly identified PSMs/peptides/proteins by individual database search algorithms in Round 1 of APIR. **B.** Proportions of combinations that show a non-negative percentage increase (green bars) in true PSMs (first column), true peptides (second column), and true proteins (third column). **C**. The indices of database search algorithms in (A) and the implementation of APIR in Round 1. Based on the benchmarking results in Figure 1C, in Round 1 of APIR, we applied p-value-free APIR-FDR to Byonic, Mascot, SEQUEST, and MS-GF+, and we applied p-value-based APIR-FDR to MaxQuant. In later rounds of APIR, we used p-value-based APIR-FDR for FDR control.

**Figure S7.**
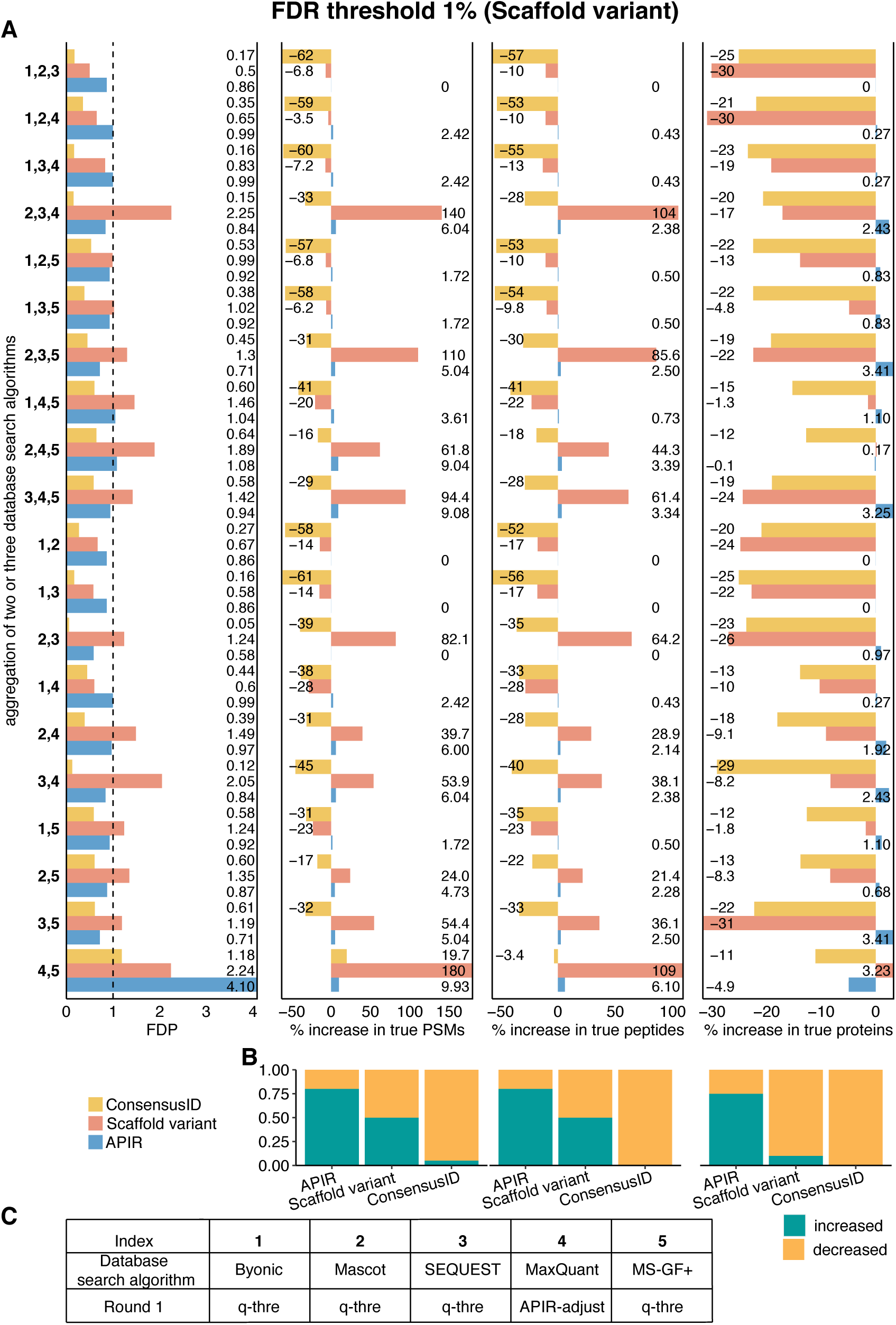
On the proteomics standard dataset, comparison of APIR, Scaffold variant, and ConsensusID at the FDR threshold *q* = 1% in terms of FDR control and power. We set Scaffold’s peptide threshold to be 1% FDR and varied its protein threshold to find the maximal number of identified peptides. **A**. FDPs (first column), the percentage increases in true PSMs (second column), the percentage increases in true peptides (third column), and the percentage increases in true proteins (fourth column) after aggregating two or three database search algorithms out of the five (Byonic, Mascot, SEQUEST, MaxQuant, and MS-GF+). The percentage increase in true PSMs/peptides/proteins is computed by treating as the baseline the maximal number of correctly identified PSMs/peptides/proteins by individual database search algorithms in Round 1 of APIR. **B**. Proportions of combinations that show a non-negative percentage increase (green bars) in true PSMs (first column), true peptides (second column), and true proteins (third column). **C**. The indices of database search algorithms in (A) and the implementation of APIR in Round 1. Based on the benchmarking results in Figure 1C, in Round 1 of APIR, we applied p-value-free APIR-FDR to Byonic, Mascot, SEQUEST, and MS-GF+, and we applied p-value-based APIR-FDR to MaxQuant. In later rounds of APIR, we used p-value-based APIR-FDR for FDR control.

**Figure S8.**
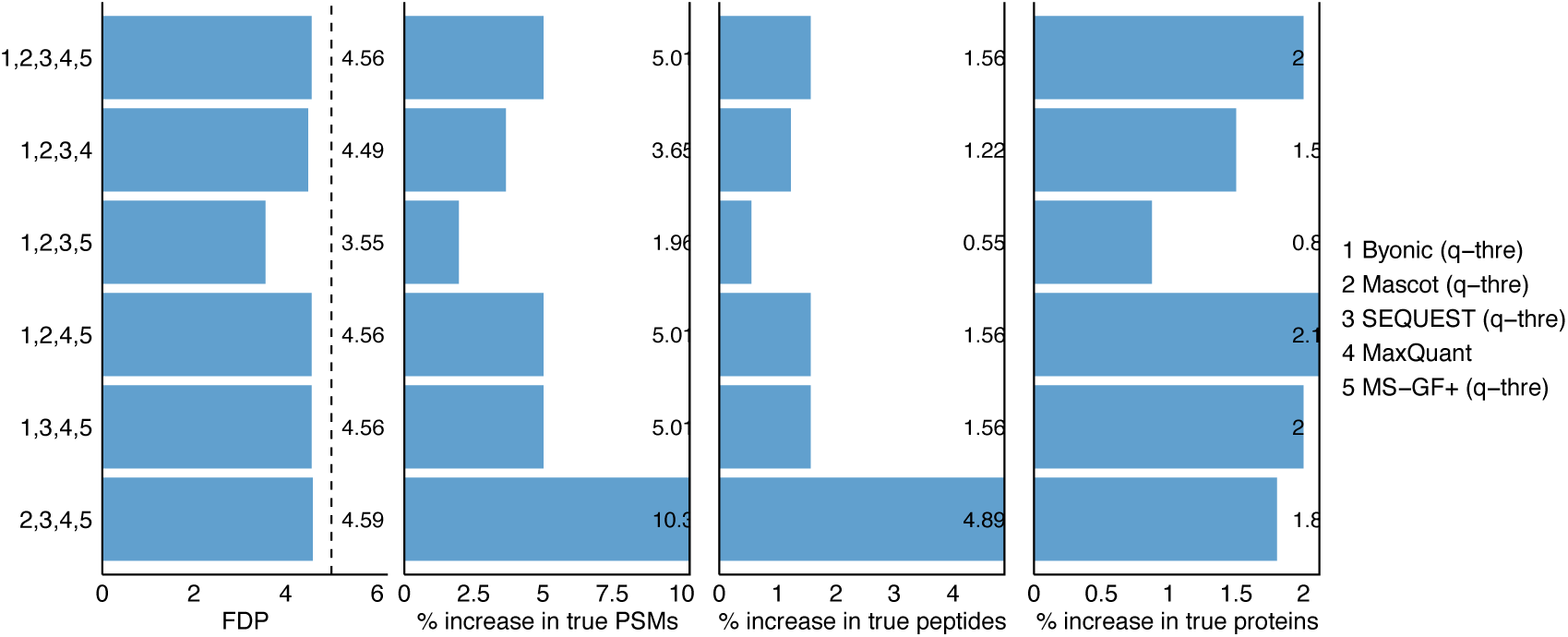
On the proteomics standard dataset, comparison of APIR and individual database search algorithms at the FDR threshold *q* = 5% in terms of FDR control and power. FDPs (first column), the percentage increases in true PSMs (second column), the percentage increases in true peptides (third column), and the percentage increases in true proteins (fourth column) after aggregating four or five database search algorithms out of the five (Byonic, Mascot, SEQUEST, MaxQuant, and MS-GF+). The percentage increase in true PSMs/peptides/proteins is computed by treating as the baseline the maximal number of correctly identified PSMs/peptides/proteins by individual database search algorithms in Round 1 of APIR.

**Figure S9.**
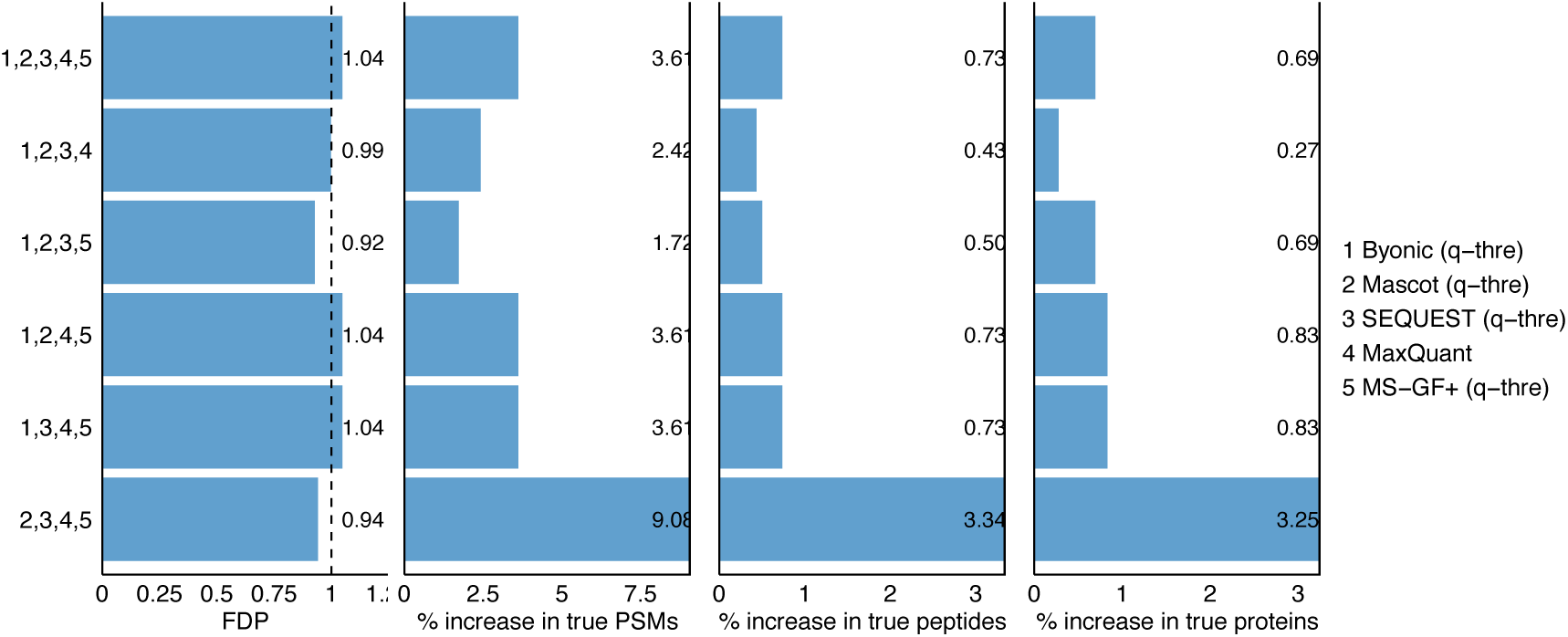
On the proteomics standard dataset, comparison of APIR and individual database search algorithms at the FDR threshold *q* = 1% in terms of FDR control and power. FDPs (first column), the percentage increases in true PSMs (second column), the percentage increases in true peptides (third column), and the percentage increases in true proteins (fourth column) after aggregating four or five database search algorithms out of the five (Byonic, Mascot, SEQUEST, MaxQuant, and MS-GF+). The percentage increase in true PSMs/peptides/proteins is computed by treating as the baseline the maximal number of correctly identified PSMs/peptides/proteins by individual database search algorithms in Round 1 of APIR.

**Figure S10.**
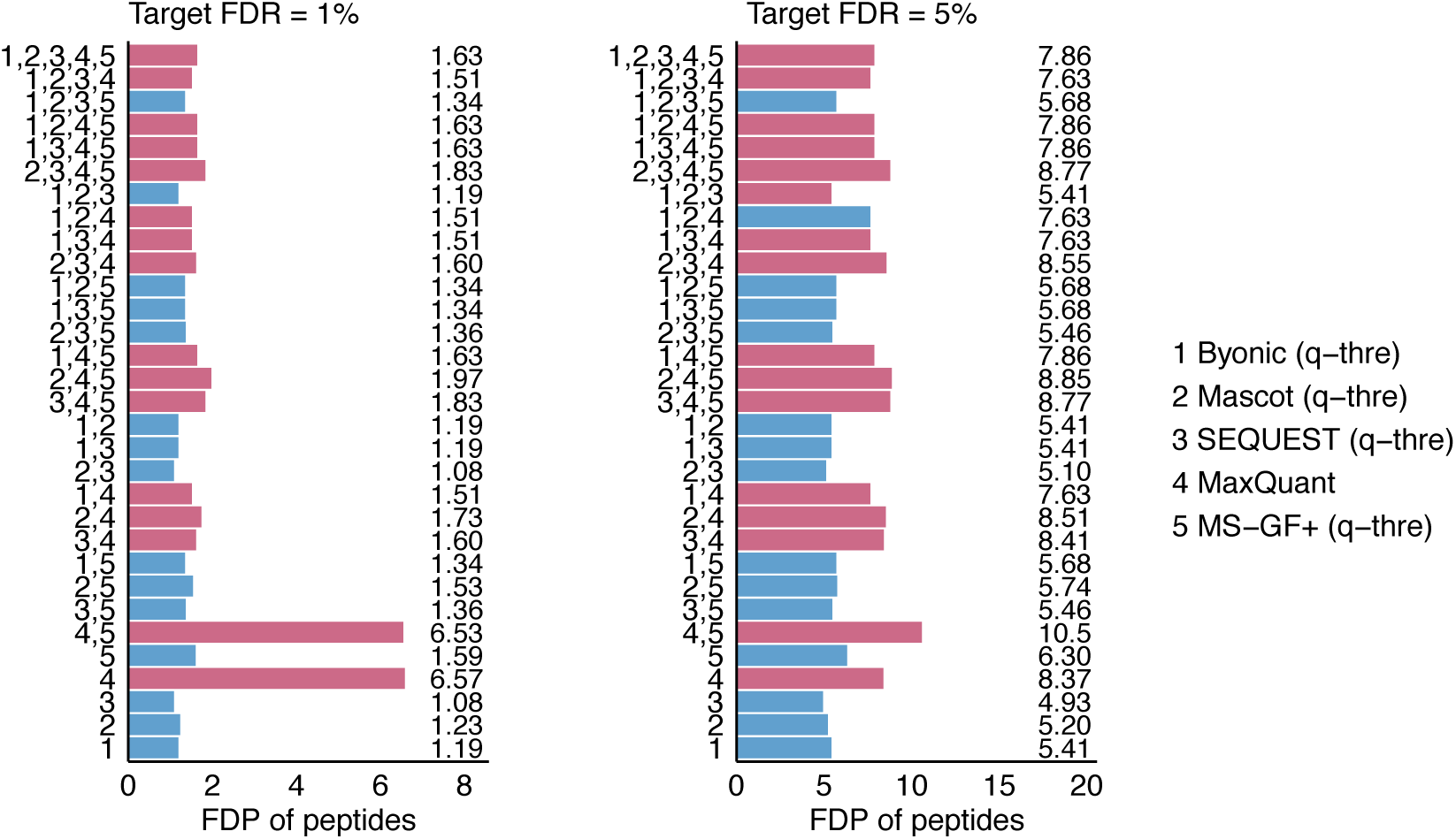
On the *Pfu* proteomics standard dataset, the FDPs of peptides identified by APIR and by individual database search algorithms at the FDR thresholds *q* = 1% (left) and *q* = 5% (right) Note that APIR only aims to control the FDR at the PSM level, not at the peptide level. Since MaxQuant on its own has a high peptide-level FDP, the algorithm combinations that involve MaxQuant are highlighted in red.

**Figure S11.**
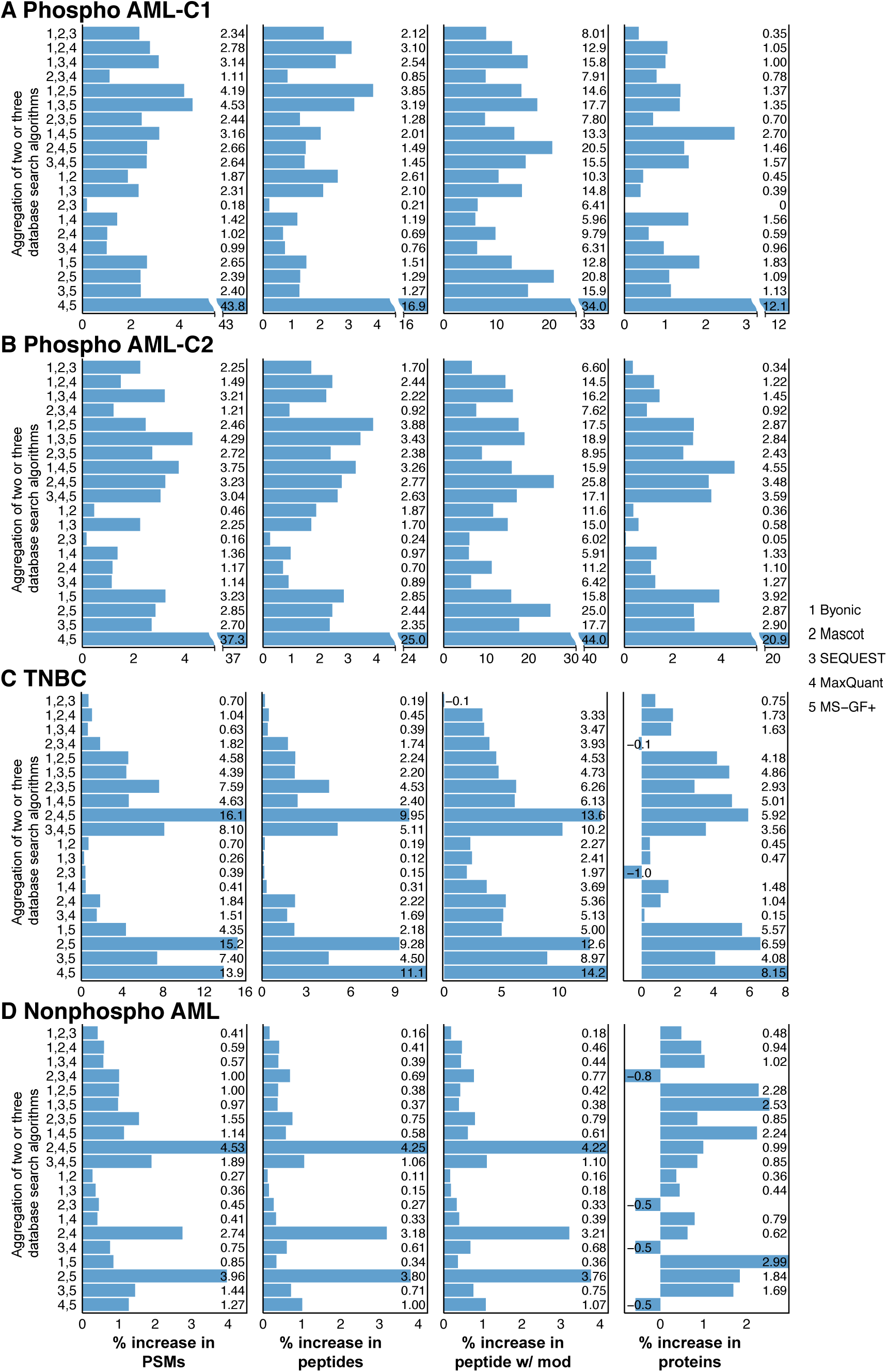
Power improvement of APIR over individual database search algorithms at the FDR threshold *q* = 1%. The percentage increases in PSMs (first column), the percentage increases in peptides (second column), the percentage increases in peptides with modifications (third column), and the percentage increases in true proteins (fourth column) of APIR after aggregating two or three database search algorithms out of the five (Byonic, Mascot, SEQUEST, MaxQuant, and MS-GF+) at the FDR threshold *q* = 1% on (**A**) the phospho AML--C1 dataset, (**B**) the phospho AML--C2 dataset, (**C**) the TNBC dataset, and (**D**) the nonphospho AML dataset. The percentage increase in PSMs/peptides/peptides with modifications/proteins is computed by treating as the baseline the maximal number of PSMs/peptides/peptides and modifications/proteins by an individual database search algorithm in Round 1 of APIR.

**Figure S12.**
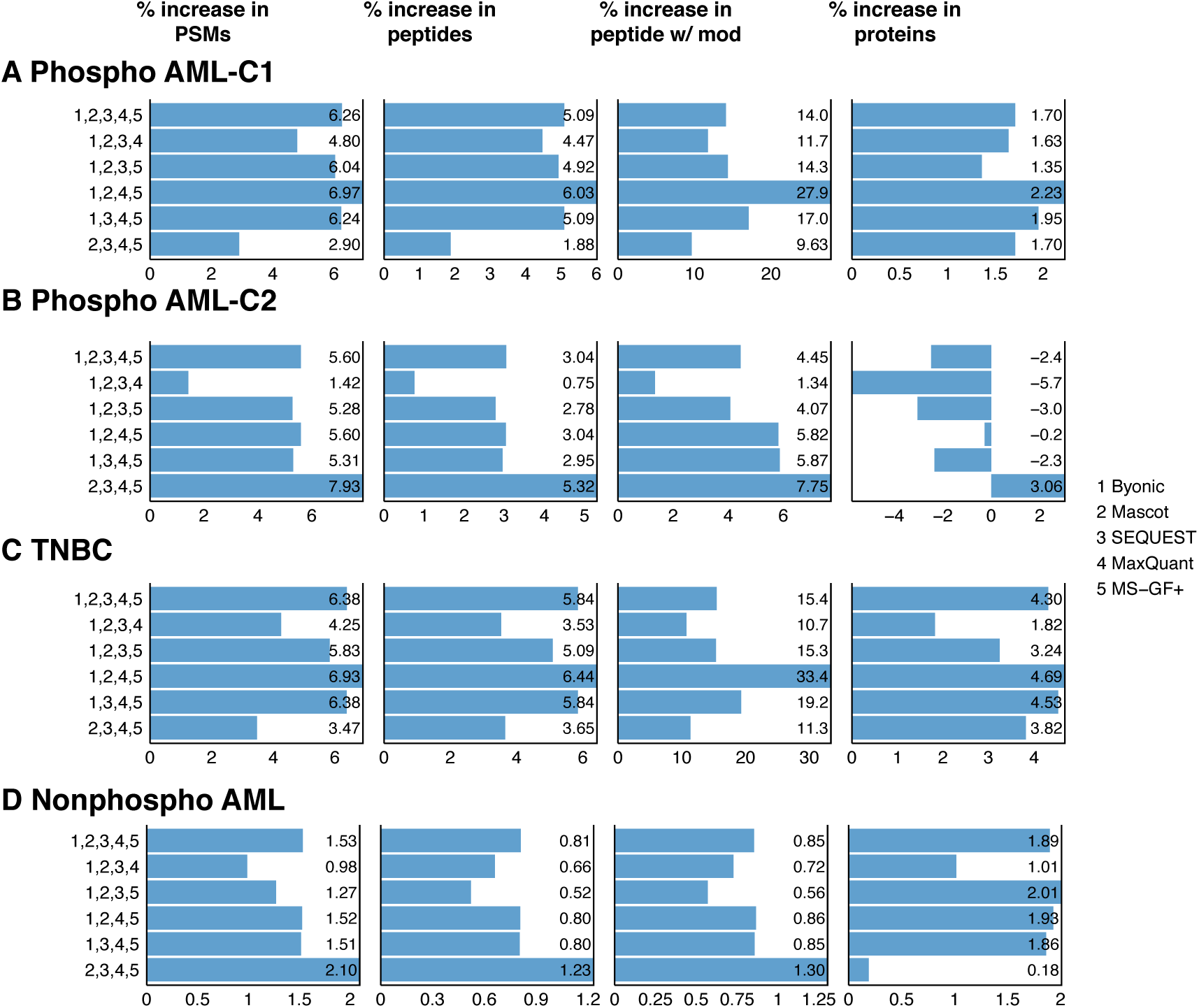
Power improvement of APIR over individual database search algorithms at the FDR threshold *q* = 5%. The percentage increases in PSMs (first column), the percentage increases in peptides (second column), the percentage increases in peptides with modifications (third column), and the percentage increases in true proteins (fourth column) of APIR after aggregating four or five database search algorithms out of the five (Byonic, Mascot, SEQUEST, MaxQuant, and MS-GF+) at the FDR threshold *q* = 5% on (**A**) the phospho AML--C1 dataset, (**B**) the phospho AML--C2 dataset, (**C**) the TNBC dataset, and (**D**) the nonphospho AML dataset. The percentage increase in PSMs/peptides/peptides with modifications/proteins is computed by treating as the baseline the maximal number of PSMs/peptides/peptides and modifications/proteins by an individual database search algorithm in Round 1 of APIR.

**Figure S13.**
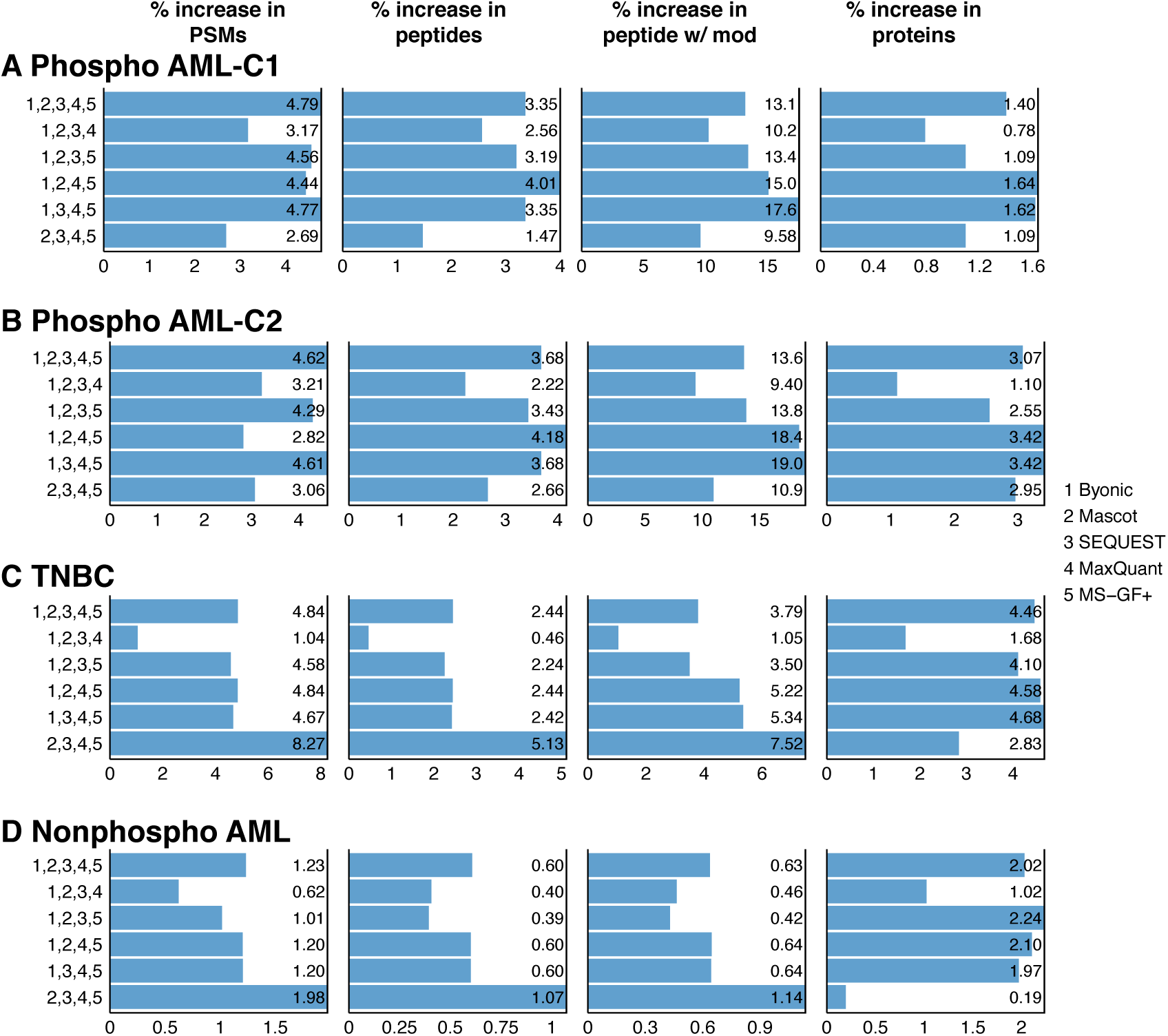
Power improvement of APIR over individual database search algorithms at the FDR threshold *q* = 1%. The percentage increases in PSMs (first column), the percentage increases in peptides (second column), the percentage increases in peptides with modifications (third column), and the percentage increases in true proteins (fourth column) of APIR after aggregating four or five database search algorithms out of the five (Byonic, Mascot, SEQUEST, MaxQuant, and MS-GF+) at the FDR threshold *q* = 1% on (**A**) the phospho AML--C1 dataset, (**B**) the phospho AML--C2 dataset, (**C**) the TNBC dataset, and (**D**) the nonphospho AML dataset. The percentage increase in PSMs/peptides/peptides with modifications/proteins is computed by treating as the baseline the maximal number of PSMs/peptides/peptides and modifications/proteins by an individual database search algorithm in Round 1 of APIR.

**Figure S14.**
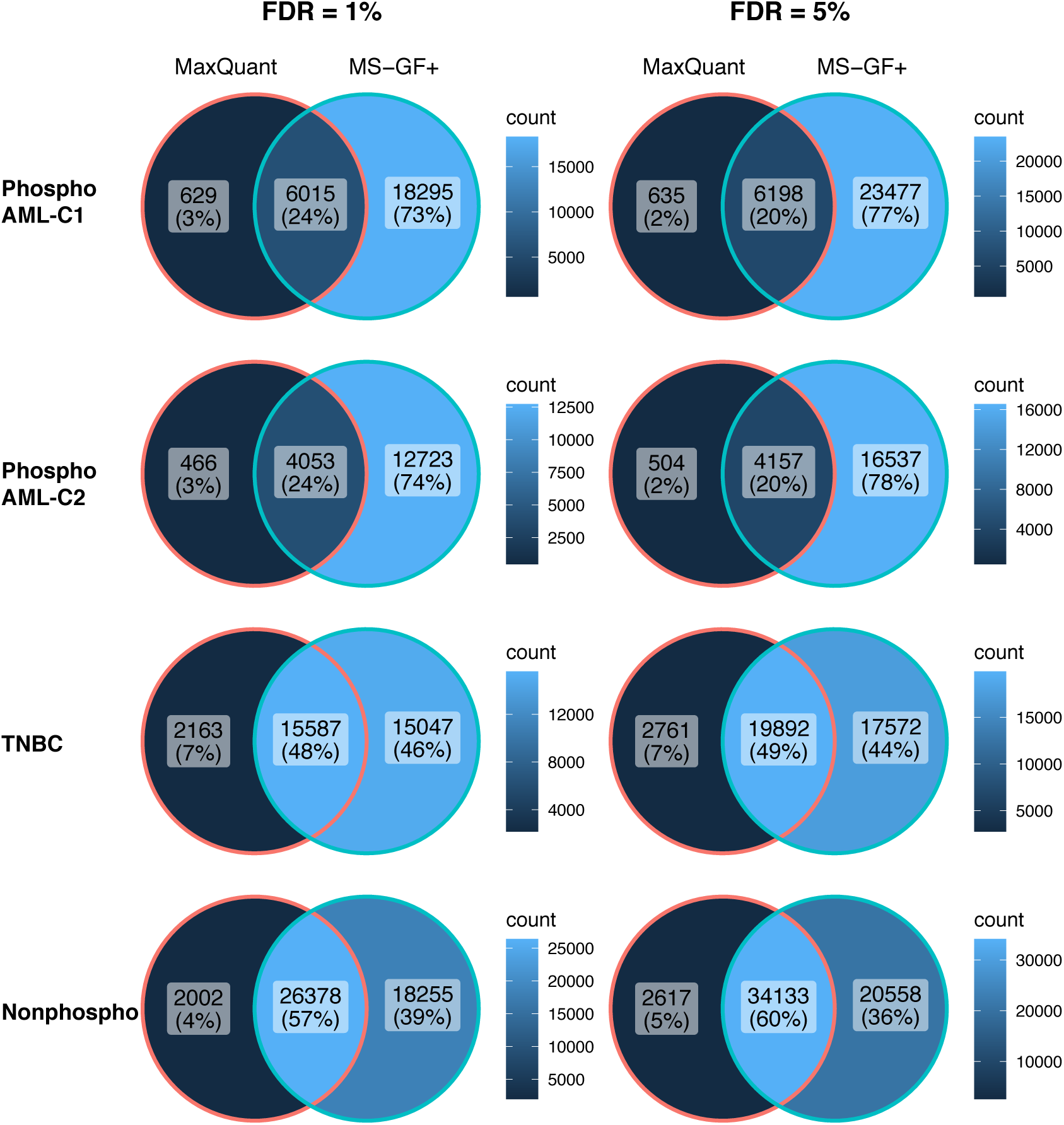
Venn diagrams of identified PSMs by MaxQuant and MS-GF+_on the four real datasets at the FDR threshold. *q* = 1% **(left) and** *q* = 5% **(right).**

**Table S1.**
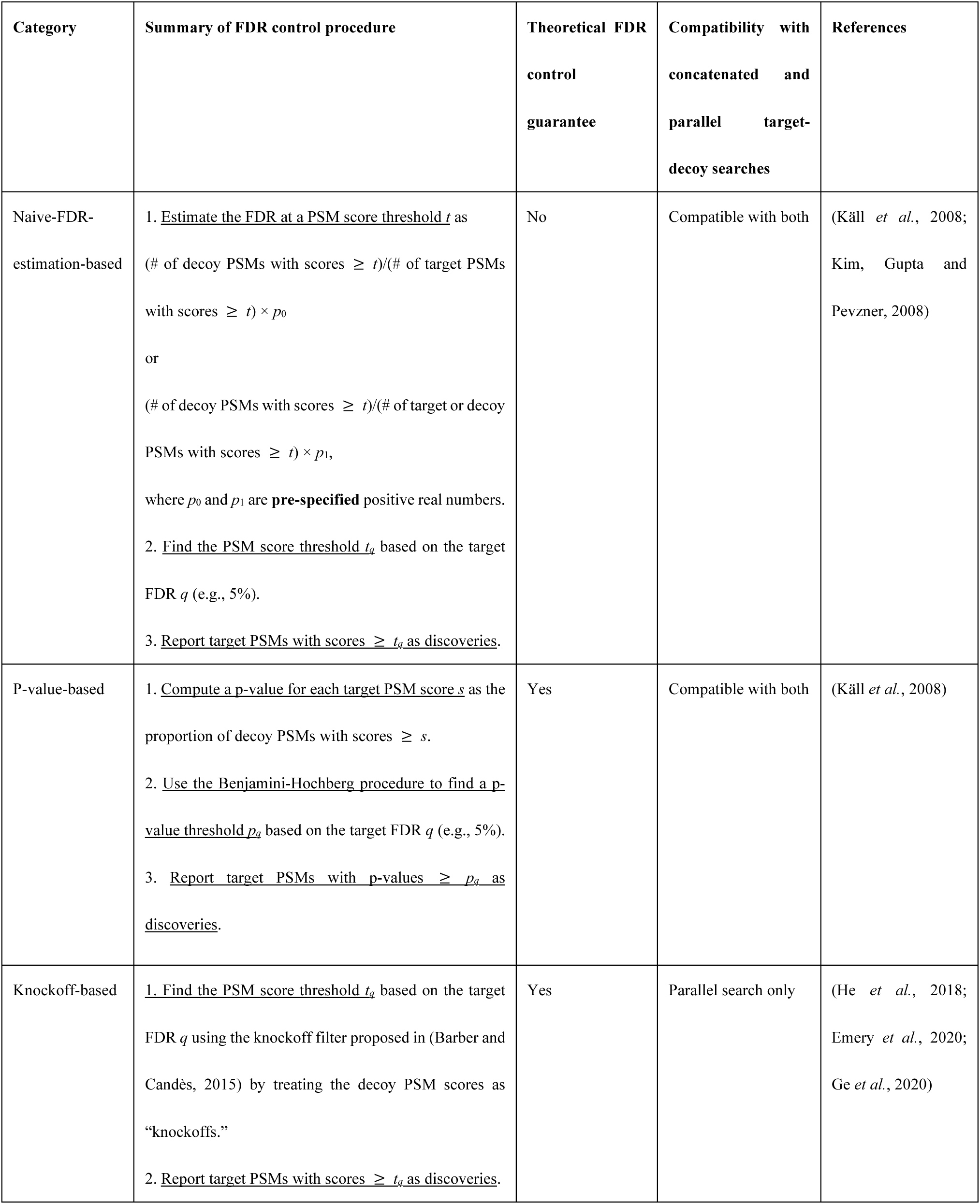
The APIR-FDR option of the five search algorithms applied on the proteomics standard dataset and other datasets (Phspho AML-C1, Phspho AML-C2, TNBC, and Nonphospho)

**Table S2.**
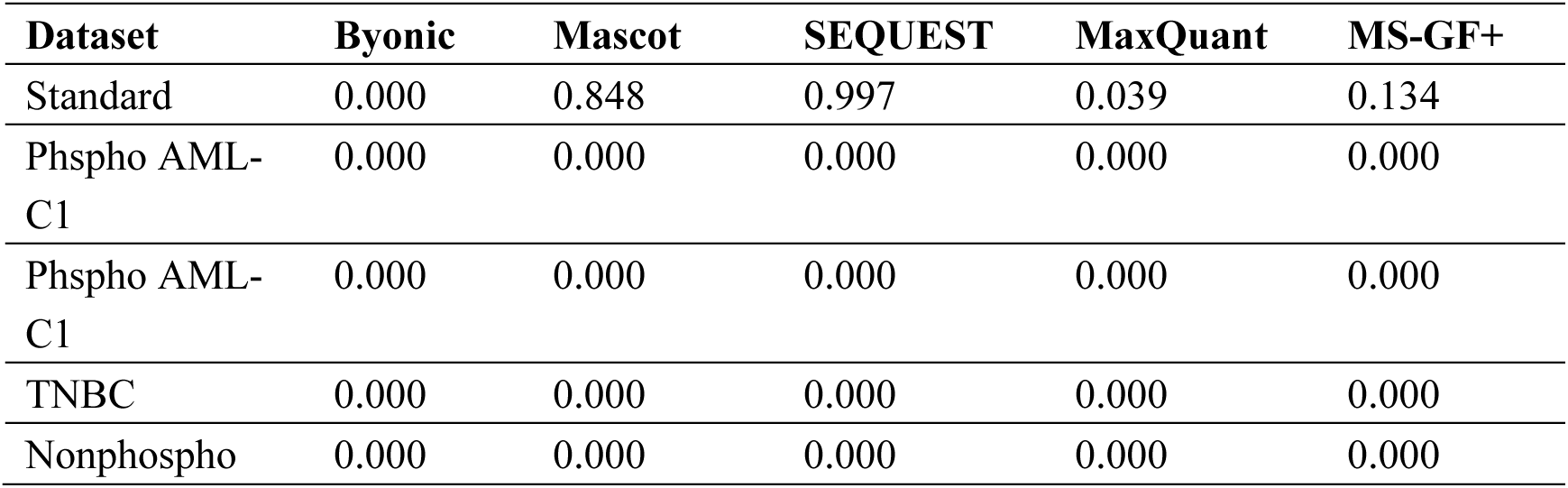
Evaluation of tandem MS spectra rescued by APIR on phospho AML-C1 and phospho AML-C2.

## Supplementary Materials

